# A conserved hub protein coordinates peptidoglycan remodeling and cell division in *Acinetobacter baumannii*

**DOI:** 10.1101/2024.09.11.612460

**Authors:** Brent W. Simpson, Amanda B. McLean, M. Stephen Trent

**Author notes:** **Author Contributions:** B.W.S. designed and performed research, analyzed data, and wrote the manuscript. A.B.M. performed research and analyzed data. M.S.T. designed research, analyzed data, and wrote the manuscript. Competing Interest Statement: Not applicable. **Classification:** BIOLOGICAL SCIENCES. Microbiology.

## Abstract

Gram-negative bacteria build a multilayered cell envelope in which the peptidoglycan layer is sandwiched between an inner membrane of glycerophospholipids and an asymmetric outer membrane composed of glycerophospholipids and lipopolysaccharide (LPS). *Acinetobacter baumannii,* however, synthesize lipooligosaccharide (LOS), an LPS variant lacking O-antigen. Although LPS/LOS is typically essential, *A. baumannii* can survive without LOS, offering the opportunity to examine how the Gram-negative envelope remains stable in the absence of this major glycolipid. We previously found that the peptidoglycan biogenesis protein NlpD, an activator of peptidoglycan degradation by cell division amidases, is critical for fitness during LOS-deficiency. Here we show that NlpD is required under these conditions because a second putative amidase activator, WthA (cell <u>w</u>all turnover <u>h</u>ub protein <u>A</u>), no longer functions in LOS-deficient cells. Mutants lacking WthA exhibited severe cell-division defects and were synthetically sick with loss of NlpD. *Acinetobacter* lack canonical periplasmic amidases, raising the question of which enzymes partner with NlpD and WthA. Previous work showed that overexpression of an *Acinetobacter* β-lactamase increased denuded peptidoglycan, a product of amidase activity. Guided by this finding, we examined the chromosomally encoded β-lactamase Oxa51 and found that its co-expression with WthA or NlpD enhanced release of amidase products, suggesting that Oxa51 participates in peptidoglycan degradation and that WthA is an amidase activator. Further, WthA influenced peptidoglycan endopeptidases and lytic transglycosylases through a network of protein interactions. Altogether, these findings identify WthA as a missing regulator in *Acinetobacter* peptidoglycan biogenesis and a hub that coordinates peptidoglycan turnover and cell division.

**Significance Statement:** Bacteria rely on a rigid cell wall, termed peptidoglycan, that must be continuously remodeled to allow growth and division. This process, known as peptidoglycan turnover, is essential for cell integrity but must be carefully controlled to prevent cell death. The critical pathogen *Acinetobacter baumannii* was missing known peptidoglycan amidases, a class of turnover enzymes, and the key activator that controls their activity during cell division. We have identified WthA as having a role in cell division most likely as an amidase activator. WthA homologs were widely distributed in bacteria and impacts two other types of turnover enzymes. We explore the possible functions of this new family of proteins that serves as a hub for impacting peptidoglycan turnover.

## Introduction

*Acinetobacter baumannii* is a Gram-negative, opportunistic pathogen with a high incidence of multi-and extreme-drug resistance (1–3). Like other Gram-negative bacteria, its resistance to many antibiotics is partly due to the structure of its dual-membrane cell envelope. The inner membrane (IM) consists of glycerophospholipids, while the outer membrane (OM) is asymmetrical with glycerophospholipids in the inner leaflet and glycolipids, either lipooligosaccharide (LOS) or lipopolysaccharide (LPS), in the outer leaflet (4, 5). Both LOS and LPS contain a core oligosaccharide attached to a negatively charged lipid A anchor, but LPS also contains a repeating O-antigen polysaccharide (6). The tight packing of these amphipathic glycolipids on the surface of the OM creates a highly impermeable barrier that contributes both to antimicrobial resistance and to the mechanical stability of the OM (7) (8–11).

Cationic antimicrobial peptides, including the polymyxin antibiotics, have taken advantage of the conserved negative charge of LPS/LOS to bind and disrupt the OM (12) (13, 14). *A. baumannii* naturally produces LOS in its OM, but in response to high levels of polymyxins can obtain mutations that completely abolish LOS synthesis (15, 16). Although this adaptation confers polymyxin resistance, it has many trade-offs including slow growth, increased sensitivity to other antibiotics, and stress in maintaining envelope integrity (11, 15–19). Few bacteria can tolerate the complete loss of LPS or LOS (15, 20–22), which makes *A. baumannii* a valuable model to explore novel mechanisms of cell envelope biology.

To identify key factors that support survival in the absence of LOS, we performed genome-wide transposon sequencing (Tn-seq) in cells with and without LOS (11). Genes impacting fitness in LOS-deficient cells were enriched for functions involved in the growth, maintenance, and division of the PG cell wall. PG is a rigid mesh with chains of alternating GlcNAc and MurNAc sugars (*N*-acetylglucosamine and *N*-acetylmuramic acid, respectively) that are crosslinked by short peptides attached to MurNAc residues (23). The PG precursor, lipid II, is synthesized in the cytoplasm and is composed of GlcNAc-MurNAc-pentapeptide subunit attached to an undecaprenyl-phosphate lipid carrier (23). Once Lipid II is flipped across the IM (24), PG synthases polymerize and crosslink new subunits into the PG layer through glycosyltransferase and transpeptidase activities, respectively (23). Because the cell wall is rigid and continuous, its expansion requires carefully regulated hydrolases (**Fig. 1A**) that cleave existing bonds to insert new subunits and to separate daughter cells after division. This remodeling process, known as PG turnover, must be precisely coordinated with synthesis to prevent uncontrolled degradation and cell lysis.

**Figure 1.**
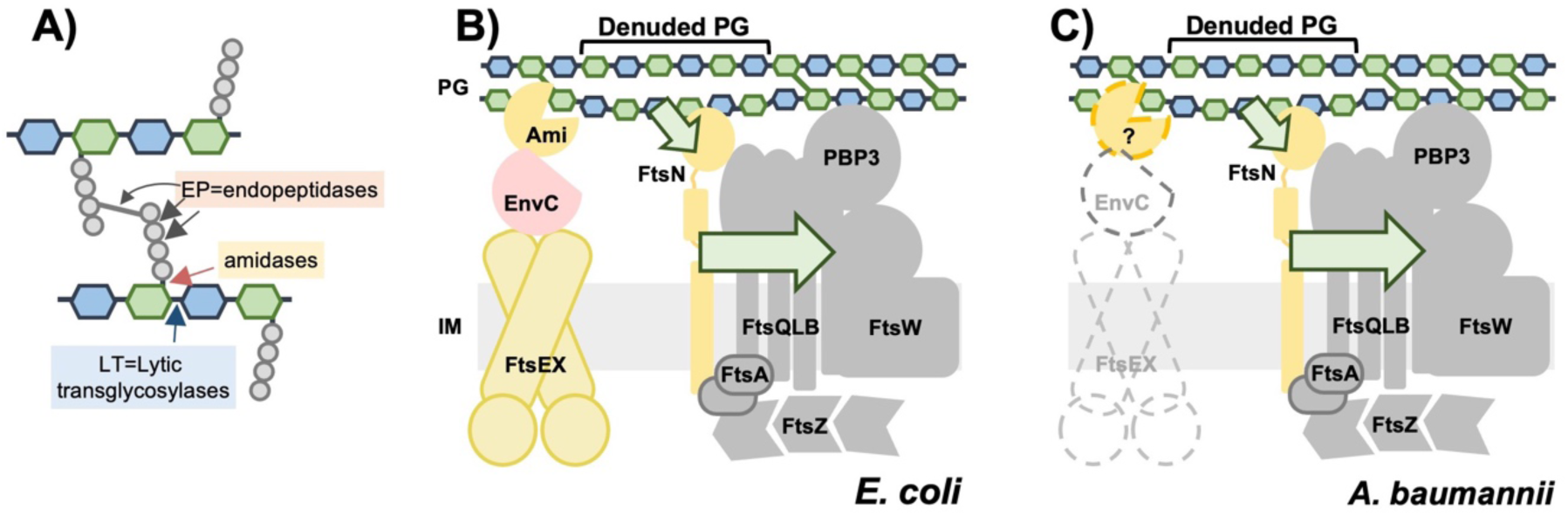
***A. baumannii* is missing homologs of key components involved in cell division.** (A) Schematic showing sites of cleavage for major classes of PG turnover enzymes. (B) In *E. coli*, amidases (AmiA, AmiB, or AmiC) that remove PG stem peptides are highly regulated. During cell division, ATP hydrolysis by FtsEX induces a conformational change in EnvC releasing a self-inhibiting helix, allowing EnvC to activate AmiA and AmiB. These amidases generate “denuded” glycan strands lacking peptide crosslinks, which are recognized by the SPOR domain of FtsN. FtsN then stimulates divisome synthases by signaling through both FtsQLB and FtsA to activate PBP3 (transpeptidase) and FtsW (glycosyltransferase). This positive feedback loop of PG synthesis and remodeling drives completion of cytokinesis. See main text for key references. (C) *A. baumannii* lacks homologs of EnvC, FtsEX, and the canonical amidases, yet retains an essential FtsN homolog, suggesting that it utilizes divergent amidases and activators to coordinate PG turnover during cell division.

Our Tn-seq screen revealed that several genes involved in cell division (*nlpD, tolA, tolB, tolQ, tolR, pal,* and *yraP*) became critical for fitness in the absence of LOS (11). Many of these genes are known to coordinate PG remodeling during cytokinesis. The Tol-Pal system, which includes TolABQR and Pal, helps couple OM constriction (25) with PG remodeling and has been implicated in coordinating cell division PG turnover through NlpD (26). YraP, an outer membrane lipoprotein of unknown function, has also been implicated in this pathway (26). NlpD itself is a degenerate member of the LytM family of PG hydrolyases, but lacks the catalytic residues for PG degradation (27). Instead, NlpD localizes to septa and activates PG amidases that cleave the amide bond between MurNAc and crosslinked peptides (27, 28). These amidases completely release crosslinking peptides and produce so-called “denuded” glycan strands that signal progression of septal PG synthesis (**Fig. 1B**). In *E. coli*, NlpD functions redundantly with other amidase activators, including EnvC and ActS (28–30). EnvC is the primary amidase activator in most proteobacteria (**Fig. 1B**), but defects in *envC* mutants are exacerbated by loss of NlpD, Tol-Pal, and YraP (26). Because loss of these genes decreased fitness in LOS-deficient *A. baumannii*, we suspected that a cell division amidase activator was no longer functioning when the OM lacks LOS. However, *A. baumannii* was missing homologs of *envC* and missing any known amidases (**Fig. 1C**).

Cell division amidases participate in a positive feedback loop that drives septal PG synthesis and promotes cell division (**Fig. 1B**). Recruitment of divisome components is highly ordered and reviewed well by others (31, 32). Important here, the FtsEX complex acts upstream to regulate EnvC (23). After the Z-ring and divisome complexes have assembled and septal PG synthesis has begun, ATP hydrolysis by FtsEX triggers a conformational change in EnvC (33) and releases an inhibitory domain from its degenerate LytM region (34–36). Activated EnvC then stimulates the amidases AmiA or AmiB to cleave peptides from the septal PG. The resulting denuded PG is recognized by the SPOR domain of FtsN (37). Binding of FtsN to these sites triggers conformational signals that travel through two independent routes: one through FtsQLB to activate PBP3/FtsW, and the other through FtsA to activate the glycosyltransferase FtsW (38–44). Increased septal PG synthesis by PBP3/FtsW accelerates recruitment of more FtsN leading to a self-enhancing cycle of septal PG synthesis that drives completion of cell division (38). *A. baumannii* contains a single, essential homolog of FtsN (45, 46), suggesting that it retains this cell division reinforcement loop (**Fig. 1C**). However, it lacks homologs of FtsEX, EnvC, and the canonical amidases AmiABC, leaving key steps in this regulatory loop unresolved.

Using the LOS-deficient background as a genetic tool, we set out to identify factors important for cell division including the missing amidase activation machinery. We discovered that HMPREF0010_01334 from strain ATCC 19606, which we designate as WthA, plays a critical role in cell division and growth. WthA is predicted to be localized to the periplasm and contains a high number of TPR (tetratricopeptide repeat) domains that facilitate protein-protein interactions. Cells lacking *wthA* exhibited severe chaining, lysis, and osmotic sensitivity. These defects were exacerbated with loss of *nlpD* suggesting that WthA functions as an amidase activator. Indeed, there is a precedent for involvement of TPR domains in amidase activation, as *Staphylococcus aureus* ActH also uses TPR domains for this purpose (47, 48). Additionally, we demonstrate that both NlpD and WthA induce formation of PG amidase products when co-expressed with the β-lactamase Oxa51, consistent with an amidase activator role. These findings also reinforce the recent proposal that *A. baumannii* β-lactamases can both degrade β-lactams and cleave PG (49, 50).

Beyond amidase activation, we find that WthA interacts with multiple PG turnover enzymes and have named it cell **<u>w</u>**all **t**urnover **<u>h</u>**ub protein **<u>A</u>**. WthA homologs are widely distributed among Proteobacteria, Bacteroidota and Chlorobiota. LbcA, an ortholog of WthA in *Pseudomonas aeruginosa*, binds to lytic transglycosylases that cleave the glycan chains of PG and is an adaptor protein for proteolysis of endopeptidases that cleave PG crosslinks (47, 48, **Fig. 1A**). We show that LbcA’s role in regulating multiple PG turnover enzymes is conserved in *A. baumannii* WthA. However, the existing literature and data presented here indicate that the *Pseudomonas* homolog likely does not serve as an amidase activator. Overall, our findings suggest that WthA/LbcA homologs are a family of hub proteins for regulation of PG turnover enzymes with distinct mechanisms.

## Results

### HMPREF0010_01334 is involved in cell division and does not function in LOS-deficient cells

We suspected that an unknown cell division amidase activator was not functioning in LOS-deficient *A. baumannii* based on the following logic. In *E. coli*, loss of NlpD, Tol-Pal, or YraP results in a synthetic sick phenotype with loss of the primary amidase activator EnvC (26, 28). In LOS-deficient *A. baumannii*, NlpD, Tol-Pal, and YraP were found to be critical for fitness (11) (**Fig. S1A**). *A. baumannii* lacks a homolog of EnvC, but retains an essential FtsN homolog (45, 46) (HMPREF0010_03200) that contains a SPOR domain for sensing denuded PG, the product of amidase activity (**Fig. 1**). This suggested that an amidase activator must exist, even though its identity was unknown. NlpD was an unlikely candidate for the primary activator, since it is not essential in LOS-containing cells and its disruption causes only rare chaining (**Fig. S1**). In agreement with this logic, LOS-deficient strains exhibit chaining defects (19) (**Fig. 2A**) and naturally increase expression of NlpD (16), possibly to compensate for reduced function of another amidase activator. If the missing activator were nonfunctional in LOS-deficient cells, the gene encoding it may have a positive fold-change in our LOS-deficient Tn-seq data, similar to all genes involved in synthesis and transport of LOS. For example, *lpx* and *lpt* genes are easily disrupted in LOS-deficient cells because they are dispensable in this genetic background (11). Among candidate genes, HMPREF0010_01334 (renamed here *wthA* for cell **<u>w</u>**all **t**urnover **<u>h</u>**ub protein **<u>A</u>**) fit these criteria. Tn-seq analysis showed low read counts (RPKMs) for insertions in *wthA* in cells with LOS and a positive-fold change in LOS-deficient cells (11) (**Fig. S1A**).

**Figure 2.**
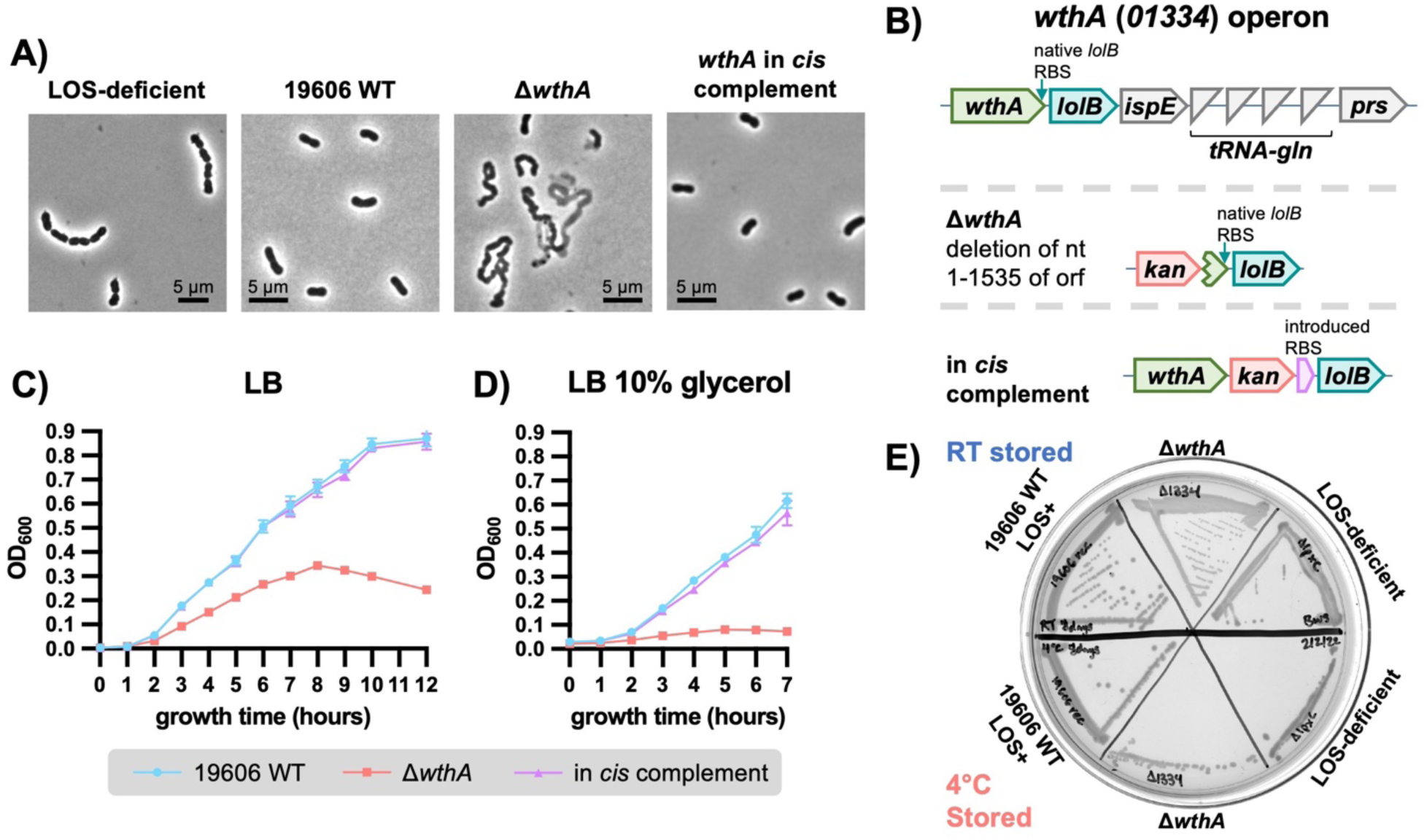
WthA (HMPREF0010_01334) impacts cell division and is nonfunctional in LOS-deficient *A. baumannii*. (A) Phase contrast microscopy of LOS-deficient and *wthA* mutants, representative of biological triplicates. (B) *wthA* operon structure and description of *wthA* mutant alleles. (C-D) Biological triplicate growth curves of *wthA* mutants in LB (C) or LB with 10% glycerol (D). Error bars are not visible when smaller than the size of the symbol. (E) Growth recovery of *wthA* and LOS-deficient mutants at 37°C after storage of colonies for 7-8 days at either room temperature (RT) or 4°C, representative of biological triplicate.

First, we constructed a *wthA* deletion mutant and assessed its morphology by microscopy. *wthA* is in an operon with *lolB* and contains the *lolB* ribosome-binding stie within its open reading frame (ORF). To prevent polar efects on *lolB,* we deleted nucleotides 1-1535 of the 1638-nucleotide *wthA* ORF (**Fig. 2B**). We also constructed an in *cis* complement strain that mimicked this alteration to *lolB*, introducing a new RBS for *lolB*, while leaving *wthA* intact (**Fig. 2B**). The *wthA* mutant exhibited a severe cell division defect characterized by long, kinked chains, evident lysis, and occasional branched cells (**Fig. 2A and S2**). The in *cis* complement strain was indistinguishable from the wild-type parent, indicating the division defects were not due to polar effects on *lolB* (**Fig. 2A**). Transmission electron microscopy confirmed these defects showing cells with detached OMs, elongated chains with several division sites, and extension of some division sites that produced the characteristic kinked appearance (**Fig. S2B**).

Next, we performed growth curves in LB and in a hyperosmotic condition, LB supplemented with 10% glycerol. In LB, the mutant showed a drastic reduction in growth rate compared to wild type or the complement (**Fig. 2D**). Under hyperosmotic conditions, Δ*wthA* cells failed to grow (**Fig. 2E**), suggesting compromised cell wall rigidity. Loss of WthA also caused cell lysis during growth, as indicated by a drop in optical density after ∼8-10 hours (**Fig. 2D**) and the accumulation of cellular debris in stationary-phase cultures (**Fig. S2A**).

To further exclude polar effects on *lolB*, we attempted genetic complementation in *trans*. The Δ*wthA* mutant was initially non-transformable, even with repeated attempts, so we disrupted a type II restriction endonuclease operon (HMPREF0010_00560 and HMPREF0010_00561) (**Fig. S3A**) to improve transformation efficiency. The resulting strain, named 19606e for <u>e</u>lectrocompetent, exhibited a ∼130-fold increase in transformation efficiency (**Fig. S3**). Complementation with the *wthA* gene alone or in combination with *lolB* in *trans* fully restored growth and morphological defects, whereas (**Fig. 3** and **S3**) a plasmid carrying only *lolB* did not (**Fig. 3** and **S3**). These results confirmed that the phenotypes of Δ*wthA* were not due to polar effects on *lolB*.

**Figure 3.**
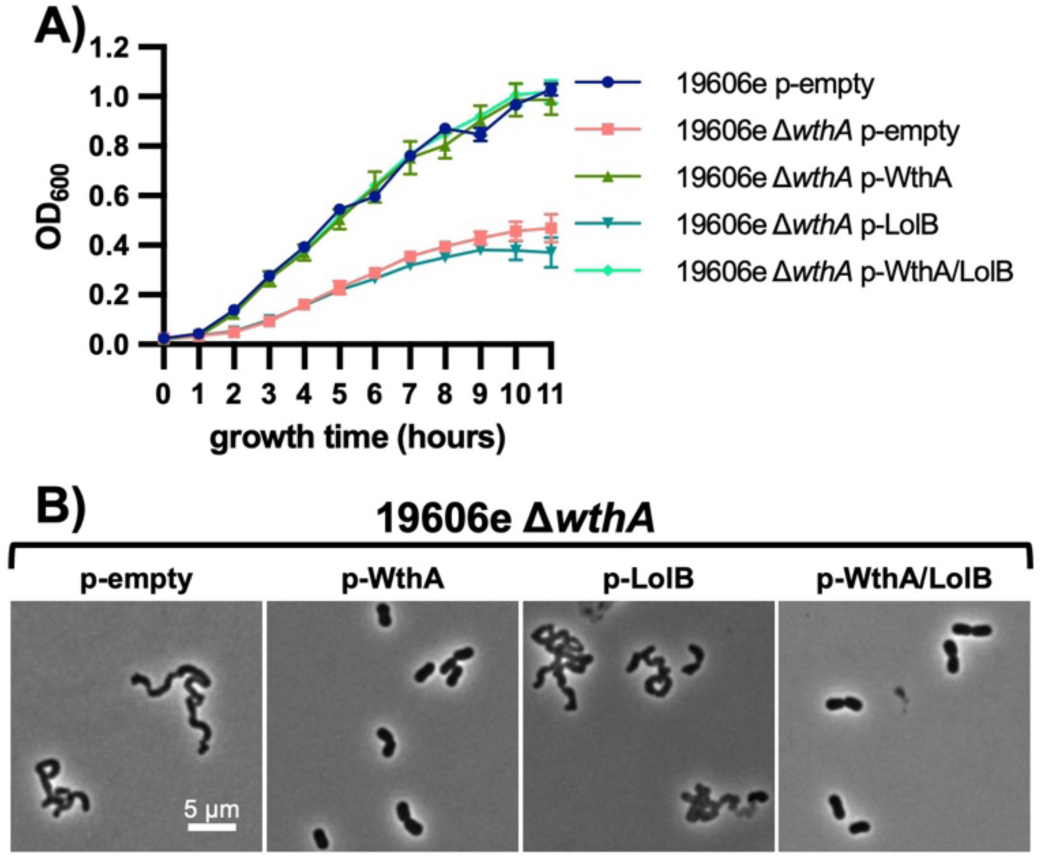
Defects from deletion of *wthA* are not due to polar effects on *lolB*. (A) Biological triplicate growth curves in LB of the 19606e Δ*wthA* mutant with different plasmids to assess for complementation. Error bars are not visible when smaller than the size of the symbol. (B) Representative phase contrast microscopy of biological triplicates of the indicated strains. Scale bar is the same for all microscopy images. (A-B) IPTG was used at a final concentration of 25 μM.

While maintaining the mutant, we noted that Δ*wthA* colonies showed poor growth after agar plates were stored at 4°C. This was striking because it is a phenotype shared by LOS-deficient strains. To test this formally, we compared the survival of Δ*wthA* (LOS^+^) and wild-type LOS-deficient cells after storage at room temperature or 4°C for 7-8 days. When colonies were restruck on LB agar and incubated at 37°C, both Δ*wthA* (LOS^+^) and LOS-deficient mutants exhibited markedly reduced growth after cold storage (**Fig. 2F**). This shared phenotype supports the idea that WthA activity is impaired in LOS-deficient cells, consistent with its proposed loss of function under these conditions.

We next attempted to characterize the cell division defect in Δ*wthA* using fluorescent labeling of the cell wall. The mutant incorporated fluorescent D-alanine derivatives poorly under all conditions tested; however, membranes and late-stage division planes could be labeled with the fluorescent FM-464 probe. The wild-type strain tolerated FM-464 labeling at all growth stages (**Fig. S4**), whereas Δ*wthA* lysed extensively when labeled during logarithmic growth (OD_600_ ∼0.5 in cuvettes). Still, cells could be labeled without impacting viability when growth began to slow (OD_600_ ∼0.8 in cuvettes, **Fig. S5-6**). Even with lysis, microscopy during rapid growth revealed numerous chains with visible late-stage division planes (**Fig. S5**). Many of these chains showed partial lysis between adjacent compartments, suggesting that septation had occurred and cytoplasmic compartments in a chain were distinct. (**Fig. S5**). With extended growth, a mixture of altered morphologies was evident including curved filaments, chains, and branched cells (**Fig. S6**), consistent with severe disruption of the division process.

### WthA impacts multiple PG turnover enzymes

The phenotypes observed were all consistent with loss of WthA causing a defect in PG biogenesis. To define which stage of PG synthesis or turnover was affected, we examined sensitivity to PG-targeting antibiotics. Loss of *wthA* did not impact sensitivity to fosfomycin, which targets MurA, suggesting that synthesis of lipid II precursors remained normal (**Table S1**). The *wthA* mutant showed increased resistance to vancomycin, which targets D-Ala-D-Ala termini, and increased sensitivity to β-lactams that inhibit penicillin binding proteins (PBPs). This pattern suggested that transpeptidation activity by PBPs was low, resulting in a build-up of uncrosslinked D-Ala-D-Ala (**Table S1**). Since amidase activity normally signals FtsN-dependent activation of the FtsW/PBP3 synthase complex (**Fig. 1B)**, this antibiotic profile supported the possibility that WthA functions as a cell division amidase activator.

Suppressor selections were performed during growth in hyperosmotic media to further understand the PG defect. The Δ*wthA* mutant was serially passaged in LB with increasing concentrations of glycerol. After transfer from media containing 5% to 10% glycerol, two populations regained growth and were subjected to whole-genome sequencing. Population 1 contained a frameshift mutation that would disrupt the MepM endopeptidase, *mepM*(*G156fs*), and population 2 contained a frameshift that would disrupt the MltD lytic transglycosylase, *mltD*(*V307fs*) (**Fig. S7A**). The *P. aeruginosa* WthA ortholog, named LbcA, binds both MepM and MltD (51, 52) (**Fig. S7B**). LbcA serves as an adaptor to direct three endopeptidases (MepM, PA1198, PA1199) to the protease CtpA (51, 52). LbcA also binds the lytic transglycosylases, MltD and RlpA, although these proteins are not protease substrates (52). The isolation of *mepM* and *mltD* suppressors in *Acinetobacter* suggested that WthA influences these enzymes through a conserved mechanism similar to *Pseudomonas* LbcA (**Fig. S7B-C)**. To test this directly, both *mepM* and *mltD* were deleted individually or in combination with *wthA*. The *mepM* and *mltD* single mutants grew normally in LB and LB with 10% glycerol (**Fig. S7D-E**). When combined with Δ*wthA*, both the Δ*mepM* Δ*wthA* and Δ*mltD* Δ*wthA* double mutants partially rescued growth in LB with 10% glycerol and fully rescued growth in LB (**Fig. 4A-B**).

**Figure 4.**
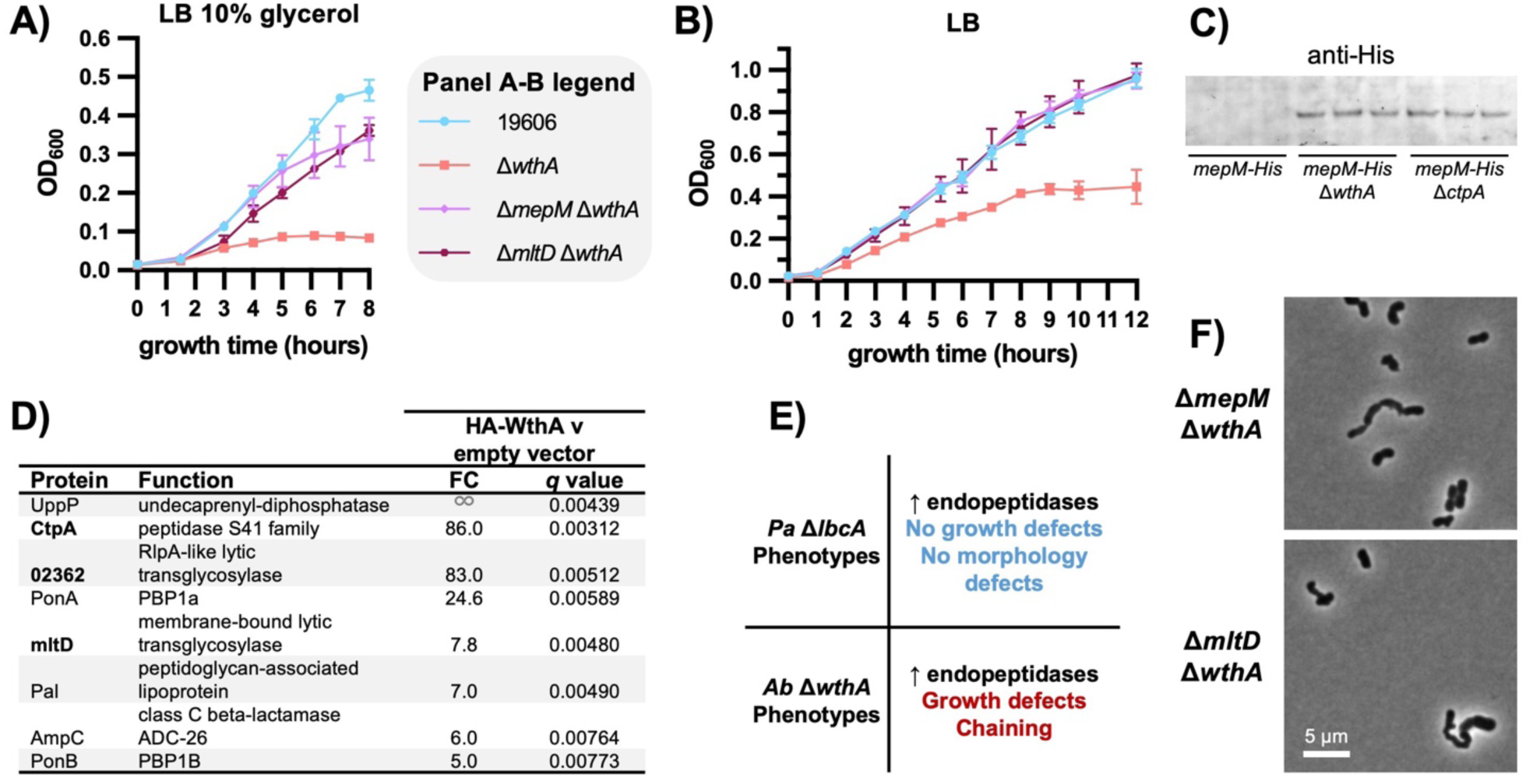
WthA is a hub protein that interacts with multiple peptidoglycan turnover enzymes. (A-B) Growth curves of Δ*wthA* suppressors in LB + 10% glycerol (A) or LB alone (B). Data are from biological triplicate and error bars are not visible when smaller than the size of the symbol. (C) Immunoblot showing chromosomally expressed MepM levels in the presences and absence of WthA and the protease CtpA. Biological triplicates are shown. (D) PG biogenesis proteins co-purified with HA-tagged WthA identified by shot gun proteomics following immunoprecipitation. FC denotes fold-change in peptide abundance relative to control immunoprecipitations from strains containing empty vector. (E) Listing of phenotypes observed for Δ*lbcA* and Δ*wthA* mutants. (F) Phase contrast microscopy of Δ*wthA* suppressors, representative of biological triplicates.

These results suggested that WthA impacts MepM and MltD similarly to LbcA. To determine whether WthA facilitates CtpA-dependent proteolysis of MepM, we monitored protein levels of a chromosomal His-tagged allele of *mepM* in strains with or without WthA and CtpA. MepM-His was undetectable when both WthA and CtpA were present but increased dramatically when either protein was absent (**Fig. 4C**), indicating that WthA participates in proteolytic regulation of MepM. To test if WthA associates with multiple PG biogenesis proteins similar to LbcA, HA-tagged WthA was overexpressed, crosslinked with EGS [ethylene glycolbis(succinimidyl succinate)], and immunoprecipitated from wild-type *A. baumannii*. The elutant was subjected to mass spectrometry to identify co-immunoprecipitated proteins. Significantly enriched hits were defined relative to two independent controls: an empty plasmid and a cytoplasmic HA-tagged control protein (ElsL-HA) (**Dataset S1**). HA-WthA co-purified with CtpA, an RlpA-like protein, and MltD, interactions conserved with its LbcA ortholog (**Fig. 4D**). Additional PG biogenesis proteins were also enriched in the pull-down (**Fig. 4D**), confirming that WthA functions as a hub for multiple PG turnover enzymes in *Acinetobacter*.

We suspected that changes in MepM and MltD function alone were unlikely to explain the pronounced cell division defects of Δ*wthA* because it is well documented that *Pseudomonas lbcA* mutants show no growth or cell division defects (**Fig. 4E)** (51). Increased activity of MepM and MltD would be expected to be highly toxic if PG synthesis were already reduced, as suggested by the antibiotic sensitivity profile of a Δ*wthA* strain (**Table S1**). A similar effect has been observed for β-lactam inhibition of PG synthesis, during which lysis is attributed to PG degradation still being high while synthesis is low (28, 53–56). The ability of Δ*mepM* and Δ*mltD* to rescue Δ*wthA* cell division defects was assessed and both Δ*mepM* Δ*wthA* and Δ*mltD* Δ*wthA* retained some chaining and morphological defects (**Fig. 4F and S7F**). This suggested that our initial suppressor mutations rescue growth by reducing PG degradation to match decreased PG synthesis. Notably, *P. aeruginosa* utilize EnvC as the primary amidase activator for cell division (57), whereas *A. baumannii* are missing EnvC (**Fig. S7B-C**). Together, these observations led us to hypothesize that WthA also contributes to amidase activation during cell division, providing a functional link between its hub activity and septal PG turnover.

### WthA and NlpD alterations have synthetic effects, suggesting partial redundancy

Since LOS-deficient cells increase expression of NlpD (16) and WthA activity is compromised in this background, we first tested whether overexpression was beneficial to *wthA* mutants. Overexpression of NlpD from an IPTG-inducible plasmid slightly rescued the growth defect of Δ*wthA* but did not correct cell chaining or morphological defects (**Fig. 5A-B and S8A**). These data are consistent with partial redundancy between NlpD and WthA as amidase activators.

**Figure 5.**
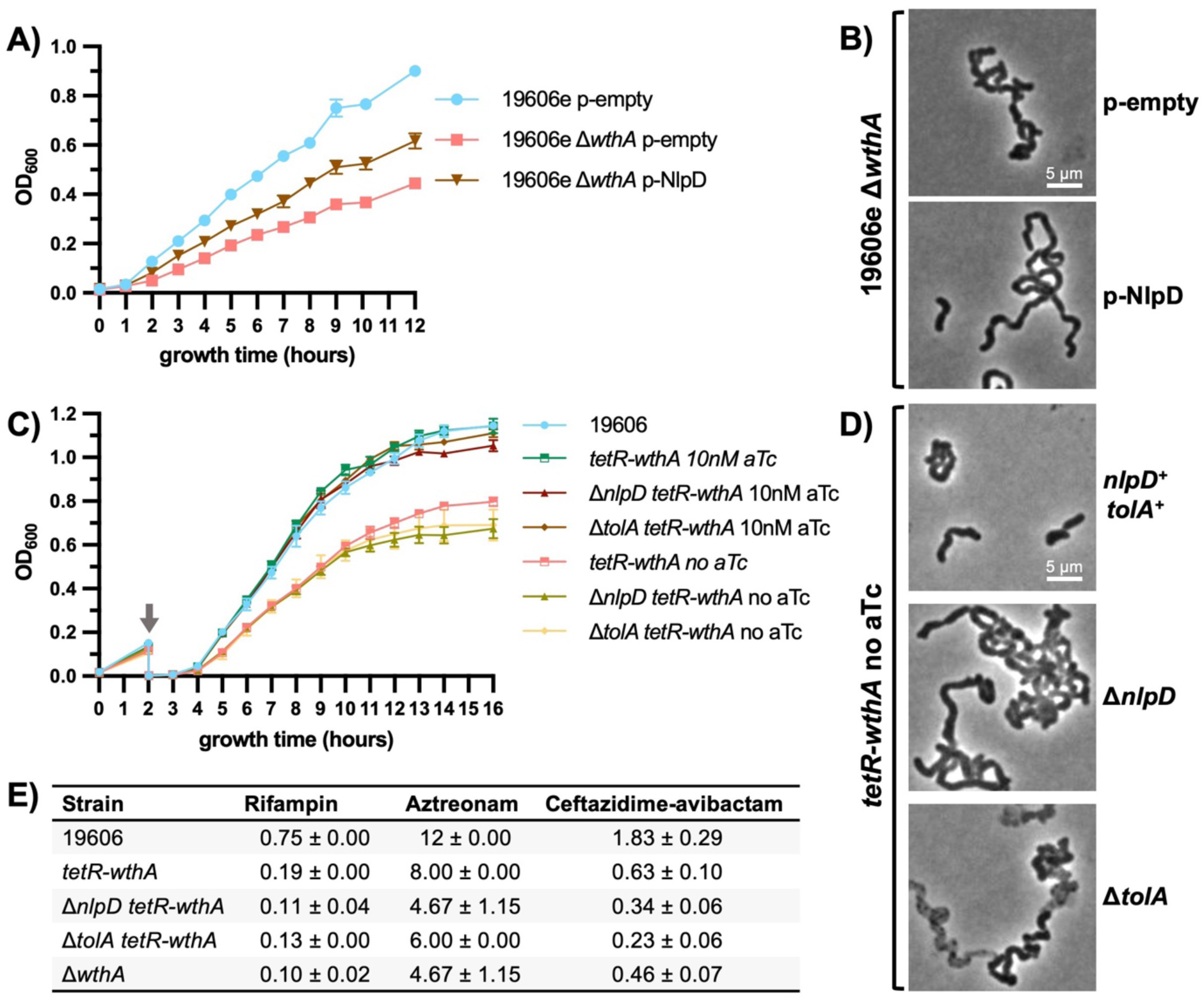
WthA and NlpD are functionally redundant, but not equivalent. (A-B) Growth curves (A) and phase-contrast microscopy (B) of the 19606e Δ*wthA* strain with or without *nlpD* overexpression. LB was supplemented with 10 µM IPTG. (C) Biological triplicate growth curves in LB with or without 10 nM aTc inducer. A single back-dilution (indicated by arrow) was used to facilitate depletion of the aTc-regulated *wthA* allele. (A,C) Error bars indicate standard deviation and are not visible when smaller than the size of the symbol. (D) Phase-contrast microscopy of *wthA* depletion strains grown in LB without aTc. (B,D) Representative of three biological replicates. (E) Minimal inhibitory concentrations of select antibiotics for *wthA* depletions strains grown without aTc. Average and standard deviation of biological triplicates are provided.

To test for possible synthetic genetic interactions, *nlpD* was deleted in a *wthA* depletion strain in which expression is controlled by an anhydro-tetracycline (aTc)-inducible promoter. We also combined Δ*tolA* with the *wthA* depletion allele to assess another pathway that impacts NlpD function. Depletion of *wthA* alone (no aTc) resulted in a growth defect and mild chaining, although less severe than the complete deletion (**Fig. 5C-D**). In contrast, deleting *nlpD* or *tolA* in combination with *wthA* depletion resulted in severe exacerbation of chaining (**Fig. 5C-D and S8B**) and only a modest further decrease in growth.

Assessment of antibiotic sensitivity supported these interactions. The *wthA* depletion strain did not have as severe of sensitivities to rifampicin, aztreonam, and ceftazidime-avibactam as compared to the deletion mutant (**Fig. 5E**). However, combining Δ*nlpD* or Δ*tolA* with the *wthA* depletion allele increased antibiotic sensitivities to levels comparable to Δ*wthA* (**Fig. 5E**). Together, these phenotypes demonstrate that loss of NlpD or Tol-Pal is synthetic sick with reduced WthA function and support a model in which WthA acts as an amidase activator that is partially redundant with NlpD.

### WthA and NlpD stimulate a toxic amidase activity when co-expressed with β-lactamase, Oxa51

*A. baumannii* lacks homologs of known amidases but retains the amidase activator NlpD, suggesting this organism may employ a different family of amidases. Recently, it was observed that overexpression of β-lactamases, including plasmid-encoded Oxa23 and chromosomally-encoded AmpC, caused physiological changes in *A. baumannii* (49, 50). Oxa23 overexpression increased amidase products, PG strands denuded of peptide crosslinks (49), while *A. baumannii* AmpC displayed weak D-Ala-D-Ala carboxypeptidase activity (50). Consistent with a possible connection, AmpC was identified among WthA interaction partners (**Fig. 4D**). Oxa23 is not present in our strain, but all *A. baumannii* strains carry a chromosomal *oxa51* allele. Oxa51 has poor carbapenemase activity compared with Oxa23 due to a sterically hindered β-lactam cleavage site (58). This conservation despite weak β-lactamase activity suggest that Oxa51 may have an alternative physiological function. Serine β-lactamases are distantly related to PG hydrolases, including endopeptidases and D-Ala-D-Ala carboxypeptidases (59), suggesting that they could act on PG peptides. In our hands, Oxa51 conferred no resistance to cephalosporins and only weak resistance to first-, third-, and fourth-generation penicillins compared with the well-characterized β-lactamase, TEM-1 (**Fig. S9**). Therefore, we examined whether Oxa51 has amidase activity that could be enhanced by NlpD or WthA.

During plasmid construction, it was immediately evident that co-expression of NlpD or WthA with Oxa51 was toxic to *E. coli* cloning strains and expression was therefore repressed during strain maintenance. When introduced into wild-type *E. coli* (W3110) or *A. baumannii*, co-expression of NlpD/Oxa51 or WthA/Oxa51 was highly toxic (**Fig. 6A-B and S10A-D**), whereas expression of the putative activators or Oxa51 alone showed no toxicity. These results suggested that NlpD and WthA may activate Oxa51 to trigger uncontrolled PG degradation. Oxa51 is expressed at low levels in *A. baumannii,* which may explain why it was not detected by mass spectrometry in the HA-WthA pull-down experiments. However, direct protein-protein interactions were obvious by reciprocal co-immunoprecipitations experiments. HA-tagged NlpD and WthA both co-purified with His-tagged Oxa51, and vice versa (**Fig. S11**).

**Figure 6.**
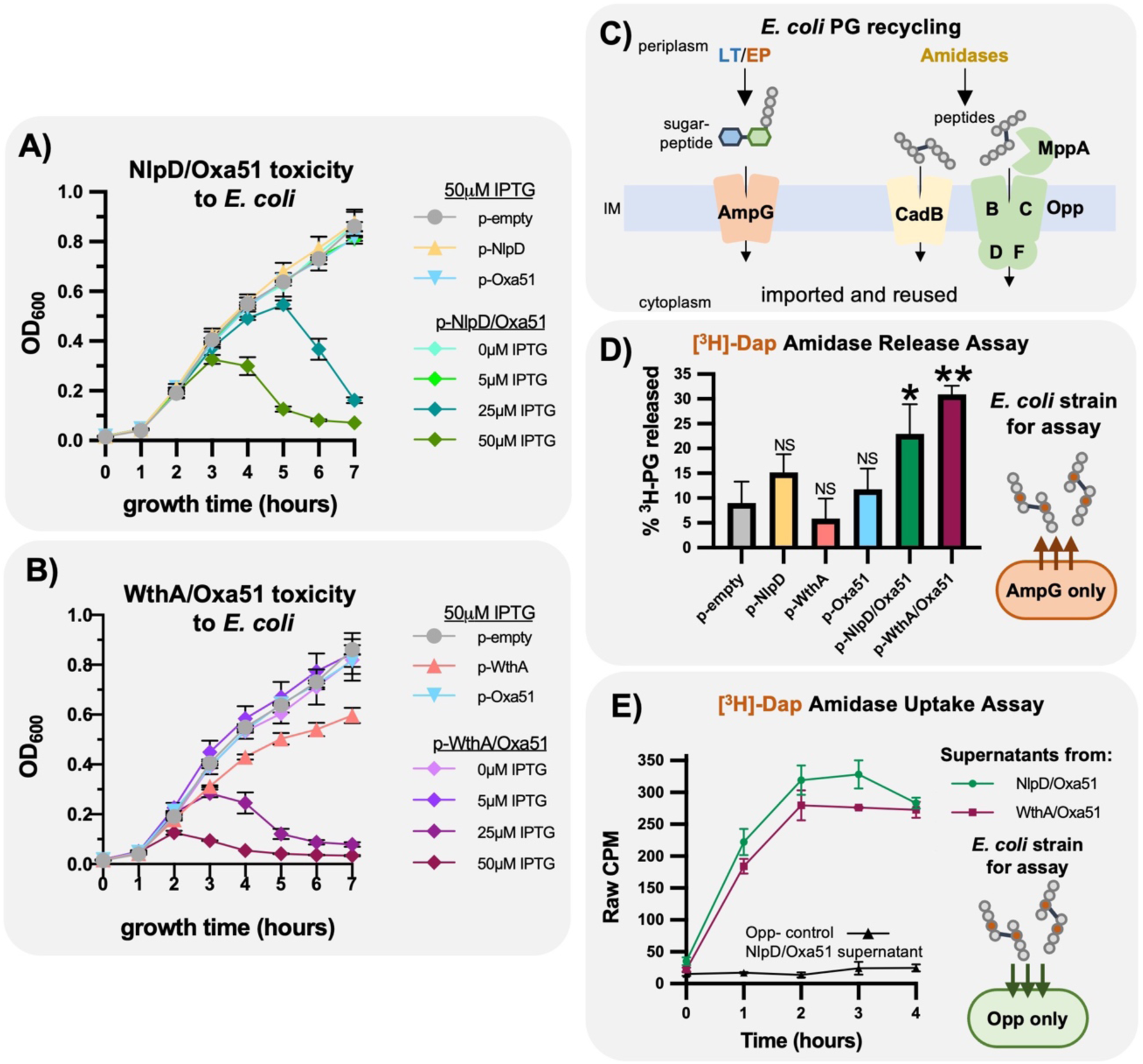
***A. baumannii* WthA and NlpD stimulate a toxic amidase activity when co-expressed with the β-lactamase Oxa51.** (A-B) Biological triplicate growth curves of *E. coli* expressing only Oxa51 or in combination with putative amidase activators, NlpD (A) or WthA (B). Concentrations of IPTG are indicated. (C) Diagram illustrating *E. coli* import systems that recycle PG turnover products. AmpG imports sugar-linked muropeptides generated by endopeptidases (EP) and lytic transglycosylases (LT). MppA/Opp and CadB import freed PG peptides released by amidases. (D) Amidase Release Assay: The AmpG-only *E. coli* strain (W3110 Δ*opp* Δ*cadB*) specifically releases amidase-freed PG peptides into the supernatant while recycling other PG fragments. Cells were radiolabeled with [^3^H]-Dap, washed to remove excess label, and release of [^3^H]-amidase products was quantified following sub-toxic expression of Oxa51 alone or in combination with NlpD or WthA (5 µM IPTG for all). (E) Amidase Uptake Assay: Spent media from the Amidase Release Assay was collected, filter-sterilized, and refreshed for carbon and nitrogen sources. Uptake of the released tritiated PG peptides was monitored using an Opp-only *E. coli* strain (W3110 ΔampG ΔamiD ΔampD, p-OppBCDF) or an Opp⁻ control strain (W3110 ΔampG ΔamiD ΔampD ΔoppABCDF, p-empty). Counts per minute (CPM) of cell pellets reflect imported radiolabel. Error bars indicate the standard deviation from three biological replicates and are not visible when smaller than the plotted symbols. Statistical significance was determined by two-tailed t-test with Welch’s correction: p < 0.05 (*), p < 0.01 (**), NS = not significant. Sub-toxic induction levels were established in **Fig. S10E–F**; controls confirming absence of lysis during the Amidase Release Assays are shown in **Fig. S10G–I**.

To directly test whether Oxa51 cleaves PG peptides, we used *E. coli* PG recycling mutants (60, 61) previously generated by our group as reporters of amidase activity (**Fig. 6C-E**). *E. coli* recycles nearly 100% of PG turnover products using specialized transporters. MppA/OppBCDF and CadB import freed peptides generated by amidases, whereas AmpG imports sugar-linked peptides produced by endopeptidases and lytic transglycosylases (60–64) (**Fig. 6C**). Therefore, a strain expressing only AmpG (Δ*opp* Δ*cadB*) will recycle sugar-linked peptides but releases freed peptides into the growth media. These freed peptides serve as a direct proxy for PG amidase activity. To measure peptide release, cells were grown in the presence of tritiated meso-diaminopimelic acid ([^3^H]-Dap) to label PG peptides. After removal of excess radiolabel, the level of [^3^H]-Dap released to the growth supernatants was quantified as a measurement of amidase activity (**Fig. 6D**). Plasmids expressing Oxa51, NlpD, and WthA alone or in combination were induced at sublethal levels (**Fig. S10E-I**) to prevent any signal from lysed cells. Expression of Oxa51, NlpD, or WthA alone produced no significant increase in released radioactivity compared to the empty vector (**Fig. 6D**). However, coexpression of NlpD/Oxa51 or WthA/Oxa51 released ∼20-30% of total cellular tritiated PG (**Fig. 6D**). Lysis was excluded by monitoring colony forming units throughout the experiment (**Fig. S10G**), and our Oxa51 coexpressions caused no detectable lysis. Although WthA/Oxa51 overexpression caused a slight growth defect, this was not due to lysis under the limited level of expression used during the assay as there was no increase in the release of DNA or ATP into growth supernatants (**Fig. S10H-I**).

Next, we verified that the released radiolabeled material consisted of freed PG peptides from amidase activity. To do this, we performed a peptide uptake assay leveraging the ability of *E. coli* Opp transport system to scavenge amidase products (60). Spent media from the amidase release assays was collected, filter-sterilized, and refreshed for carbon and nitrogen sources. These supernatants containing the released tritiated PG fragments were fed to an Opp-only strain (Δ*ampG* Δ*ampD* Δ*amiD* p-OppBCDF). This strain was blocked at several stages from being able to take in other PG fragments [see methods and ref. (60)], assuring that only Opp uptake of amidase products could occur. The Opp-only strain efficiently imported the radioactive PG products released during NlpD/Oxa51 or WthA/Oxa51 expression (**Fig. 6E**) in an Opp-dependent manner, strongly supporting that the released tritiated products were amidase-derived PG peptides.

Together, these data demonstrate that NlpD and WthA function as amidase activators that stimulate an amidase-like activity associated with Oxa51. Whether Oxa51 itself acts as the catalytic amidase or as part of a larger complex remains to be determined. Deletion of *oxa51* alone did not cause cell-division defects (**Fig. S9**), however this is consistent with the redundancy typical of amidases in other bacteria making identification of such enzymes difficult. We are in the process of identifying other potential *Acinetobacter* amidases.

### WthA/LbcA homologs are conserved in Gram-negative bacteria

All major cell division amidase activators described to date contain degenerate LytM domains, including EnvC (27, 65), DipM of *Caulobacter crescentus* (66–71), and NlpD/ActS (27, 29, 65) (**Fig. 7A**). WthA is the first putative cell division amidase activator composed instead of TPR domains. Although *S. aureus* utilizes the TPR-containing protein ActH to activate an amidase (**Fig. 7A)**, its cognate amidase cleaves uncrosslinked PG peptides to stimulate PG synthesis through a distinct mechanism that remains under investigation (47, 48). To explore whether the function of WthA might be conserved across Gram-negative bacteria, we examined the genomic distribution of *wthA* homologs. The STRING database (72) revealed that *wthA* had synteny with *lolB* that was widespread in γ-and β-Proteobacteria as well as Bacteroidota. We extended this analysis across related phylogenetic groups and identified likely *wthA* homologs in α-, β-, and γ-Proteobacteria, as well as Chlorobiota (**Fig. 7B**). Notably, several β-, and γ-Proteobacteria lacking EnvC also contained *wthA* homologs, suggesting that WthA may substitute for EnvC-type activators in these species (**Fig. 7B**).

**Figure 7.**
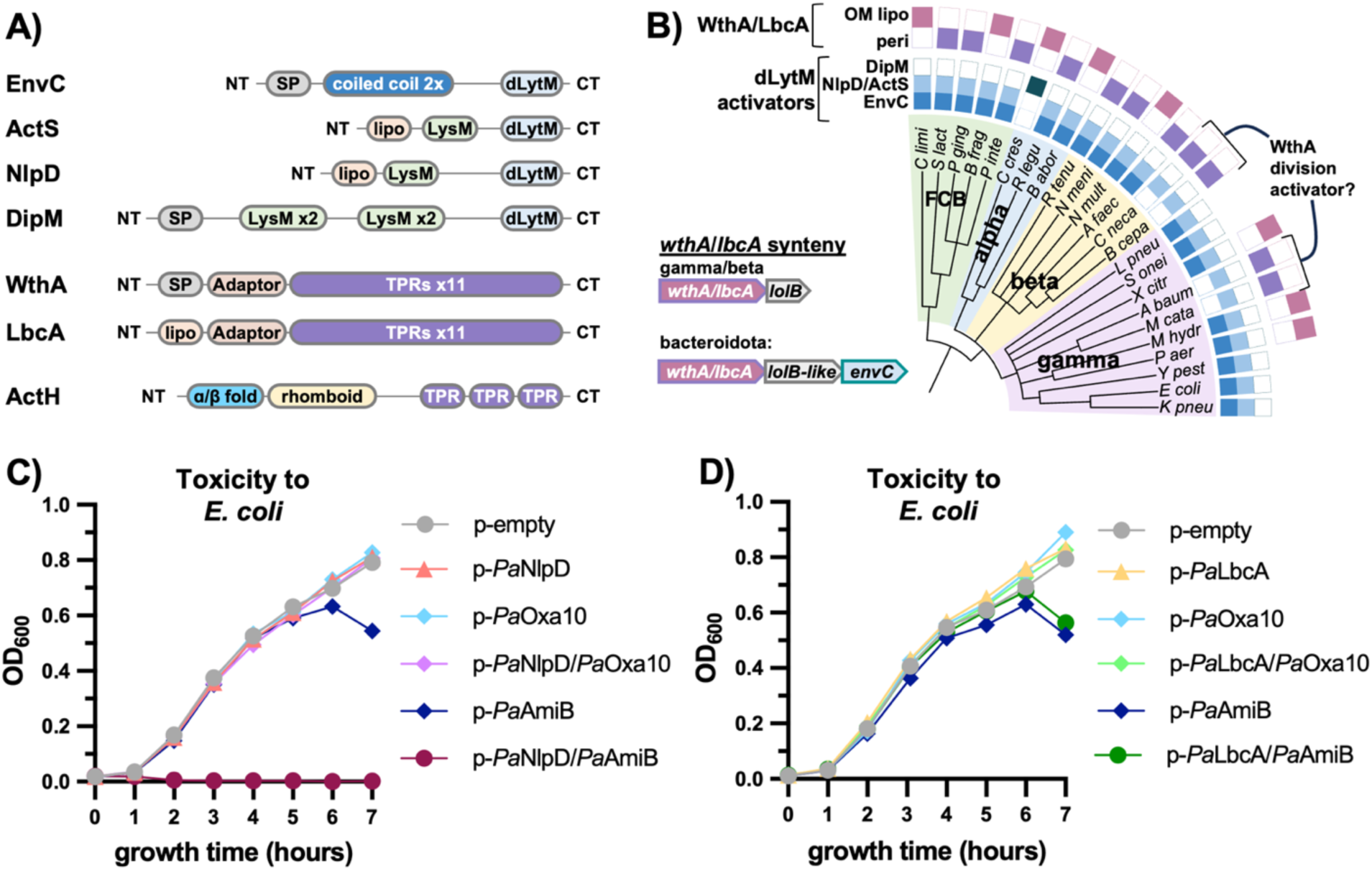
WthA/LbcA homologs are a widely conserved but differ in their mechanisms of regulating PG turnover enzymes. (A) Domain architectures of known and putative amidase activators. Degenerate LytM (dLytM) domains lacking catalytic residues are used by EnvC, NlpD, ActS, and DipM to activate cell-division amidases. In contrast, ActH and likely WthA employ tetratricopeptide repeat (TPR) domains for amidase activation. “SP” denotes periplasmic signal peptides; “Lipo” indicates lipoprotein signal peptides targeting outer-membrane (OM) anchoring. (B) Conservation of dLytM-domain activators and WthA/LbcA homologs across α-, β-, and γ-Proteobacteria and FCB (Fibrobacterota, Chlorobiota, and Bacteroidota) phyla. Predicted synteny with *lolB-like* genes is shown; these encode OM lipoproteins of unknown function that STRING analysis suggests are divergent *lolB* homologs. WthA/LbcA homologs predicted to be OM-anchored lipoproteins are marked as “OM Lipo,” whereas those predicted as soluble periplasmic proteins are labeled “Peri.” Species names and locus tags are listed in **Dataset S1.** (C-D) Growth curves of *E. coli* W3110 expressing the *P. aeruginosa* amidase, *Pa*AmiB, or the β-lactamase, *Pa*Oxa10, alone or in combination with *P. aeruginosa* NlpD, *Pa*NlpD (C) or *Pa*LbcA (D). Growth curves are representative of biological triplicates and were performed in LB with 50 µM IPTG.

*Bacteroides* species are generally thought to lack *lolB* homologs. However, in Bacteroidota we found *wthA* in synteny with OM lipoproteins genes of unknown function that STRING annotated as possible *lolB* homologs (**Fig. 7B**). These genes could encode divergent LolB proteins [20.6% identity to *E. coli* LolB by Clustal Omega alignment (73)]. Interestingly, in Bacteroidota, *envC* was located within the *wthA* operon (**Fig. 7B**), suggesting coordination of PG turnover regulators in this lineage.

Since *A. baumannii wthA* and *P. aeruginosa lbcA* mutants have distinct phenotypes, homologs of WthA/LbcA may differ in what types of PG turnover enzymes they modulate (**Fig. S7B-C**). We wondered if *P. aeruginosa* LbcA could also act as an amidase activator, as this could have been missed due to its potential redundancy with EnvC and NlpD. We cloned codon-optimized alleles of *P. aeruginosa nlpD* and *lbcA* on to plasmids along with the known amidase gene, *amiB*, and intrinsic β-lactamase, *oxa10*, referred to as *Pa*NlpD, *Pa*LbcA, *Pa*AmiB and *Pa*Oxa10, respectively. Expression of each combination of activator/amidase was evaluated for toxicity in *E. coli* serving as a proxy for amidase activation (**Fig. 7C-D**). As expected, expression of *Pa*AmiB alone caused mild toxicity given that *E. coli* NlpD and EnvC can activate amidases of this family. In agreement with existing literature showing that *Pa*NlpD functions with *Pa*AmiB (57), coexpression of these two proteins was highly toxic and allowed for no growth (**Fig. 7C** and **S12**). In contrast, *Pa*NlpD coexpression with *Pa*Oxa10 showed no evidence of toxicity (**Fig. 7C**). Coexpression of *Pa*LbcA with either *Pa*AmiB or *Pa*Oxa10 also failed to enhance toxicity (**Fig. 7D** and **S12**), indicating that LbcA does not stimulate any PG hydrolytic activities under these conditions. It remains possible that we did not test the correct amidase partner and that LbcA could activate *P. aeruginosa* AmiA or another β-lactamase. However, LbcA protein-protein interactions have been highly explored and identify AmiB as its only amidase-binding partner (52).

Since *Pa*Oxa10 had no evidence of toxicity, we wondered if *A. baumannii* Oxa51 had evolved this new activity. Exploring this, we found that Oxa51 was toxic to *E. coli* when coexpresssed with PaNlpD (**Fig. S12E**), suggesting that Oxa51 can interact functionally with NlpD from multiple species. However, PaLbcA again did not enhance toxicity when coexpressed with Oxa51 (**Fig. S12F**). These results highlight functional divergence between WthA and LbcA homologs, with WthA possibly retaining amidase-activator properties that are absent in LbcA.

## Discussion

We have provided evidence that WthA plays a critical role in cell division in *A. baumannii*. Mutants lacking WthA grew poorly and exhibited severe morphological abnormalities, including chaining during active growth (**Fig. S5**) and filamentation and branching as growth slowed (**Fig. S6**). This explains why transposon insertions in *wthA* were underrepresented in our LOS-containing transposon mutant libraries (11). The connection between WthA and cell division emerged only because its function was already impaired in LOS-deficient cells, allowing us to obtain transposon insertions and a distinct Tn-seq signature. In support of WthA dysfunction in the absence of LOS, both LOS-deficient and *wthA* mutants displayed similar cell division defects and were difficult to culture after storage at 4°C. Why WthA does not function in the absence of LOS remains unclear, but WthA may rely on an OM partner that fails to fold properly when LOS is absent.

Reduced function of WthA in LOS-deficient cells also provides a clear explanation for why NlpD, Tol-Pal, and YraP become essential for fitness under these conditions. Overexpression of NlpD partially restored growth of the *wthA* mutant, whereas loss of *nlpD* greatly intensified its division defects, consistent with the two proteins acting in overlapping pathways. These genetic relationships strongly suggest that WthA serves as an amidase activator during cell division. Notably, *A. baumannii* retains an essential SPOR-domain protein homologous to FtsN even though it lacks FtsEX and EnvC, indicating that amidase activation remains a key step in the septal synthesis feedback loop. WthA likely fulfills the role of EnvC in this system, linking PG cleavage to activation of the divisome and sustaining the self-reinforcing cycle of PG synthesis during cytokinesis. Although *A. baumannii* lacks recognizable homologs of the canonical periplasmic amidases, the conservation of three SPOR-domain proteins—one of which, HMPREF0010_03200 (FtsN), is essential—supports that this bacterium still depends on amidase-mediated PG remodeling.

Recently, overexpression of an *Acinetobacter* class D β-lactamase, Oxa23, was shown to increase denuded PG, raising the intriguing possibility that certain β-lactamases have been co-opted to perform amidase-like functions (49). We explored this hypothesis further and examined if the broadly conserved and chromosomally encoded class D β-lactamase Oxa51 could be one of the missing *Acinetobacter* amidases. Oxa51 possesses a sterically hindered β-lactamase active site that reduces its carbapenemase activity (58). Indeed, we found Oxa51 to be a weak penicillinase compared to other β-lactamases (**Fig. S9A**) suggesting functional divergence away from β-lactamases. This observation fits with the evolutionary link between class D β-lactamases and high molecular weight penicillin binding proteins (PBPs), the PG synthases from which they likely evolved (59). Notably, a chromosomally encoded class C β-lactamase, *Acinetobacter* AmpC, was recently found to possess weak D-Ala-D-Ala carboxypeptidase activity (50), supporting the broader concept that repurposed or promiscuous β-lactamases may contribute to PG metabolism in *Acinetobacter*. Oxa51 is not essential, and its deletion did not affect growth or cellular morphology (**Fig. S9**), indicating that if it is an amidase, its function is likely redundant.

Our first indication that Oxa51 could act as a PG amidase was that its expression was toxic, but only when co-expressed with NlpD or WthA. Uncontrolled activation of PG amidases is known to be highly lethal to bacteria (27), consistent with Oxa51 contributing to PG hydrolysis. To evaluate this activity directly, we leveraged *E. coli* PG recycling mutants that selectively release or import PG amidase products (60, 61). Using these strains, we designed complementary assays to measure both amidase product release and uptake. Co-expression of WthA/Oxa51 or NlpD/Oxa51 both resulted in release of PG amidase products, supporting the most likely hypothesis that WthA/Oxa51 and NlpD/Oxa51 constitute amidase and activator pairs. However, since our assays were performed in whole cells, we cannot exclude the possibility that the increased amidase activity was indirect. Further characterization will be needed to identify additional *A. baumannii* amidases and establish the mechanism of catalysis through biochemical characterization. Interestingly, the amidase activity observed when Oxa23 was overexpressed did not require the catalytic serine of the β-lactamase active site (49). This raises the possibility that class D β-lactamases, like Oxa51, employ distinct residues or alternative catalytic centers to mediate PG cleavage, representing a repurposing of their ancestral penicillin-binding protein fold.

WthA’s protein interaction profile suggested that it functions as a hub protein similar to its Pseudomonas homolog LbcA. Our data show that in *A. baumannii*, WthA: **(i)** directs CtpA-dependent proteolytic regulation of the endopeptidase MepM, **(ii)** interacts with the lytic transglycosylases RlpA and MltD, **(iii)** influences cell division, potentially as an amidase activator, and **(iv)** associates with additional PG biogenesis proteins (**Fig. 8**). The first two functions are conserved between WthA and LbcA (51, 52). However, if and how LbcA impacts lytic transglycosylases has remained elusive in *Pseudomonas,* since these enzymes are not proteolytically regulated.

**Figure 8.**
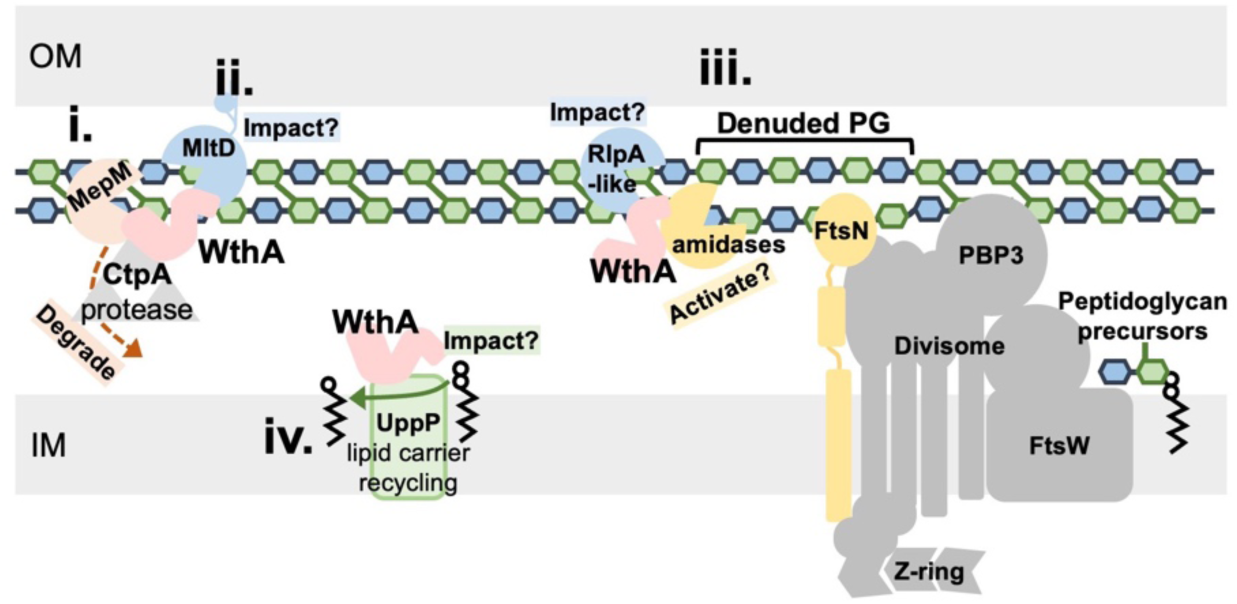
WthA functions as a central hub coordinating PG turnover in *A. baumannii*. Protein interactions and experimental evidence presented here suggest that WthA impacts multiple classes of PG turnover enzymes: **(i)**. **Endopeptidase regulation:** WthA promotes CtpA-dependent proteolysis of the endopeptidase MepM. **(ii)**. **Lytic transglycosylase control:** Suppressor analysis of Δ*wthA* disrupted MltD to rescue growth, suggesting WthA negatively regulates its activitiy to maintain PG homeostasis. The same is true for MepM supporting endopeptidase regulation. **(iii) Amidase activation**: Loss of WthA results in cell division defects that are exacerbated with alterations to NlpD. Furthermore, like NlpD, coexpression of WthA with Oxa51 was toxic and released amidase products, consistent with the protein functioning as an amidase activator. Oxa51 may represent one of the missing amidases in *A. baumannii*. Immunoprecipitations studies showed that WthA also interacts with HMPREF0010_02362, an RlpA-like lytic transglycosylase, which may act downstream of amidases to remove denuded PG strands. **(iv). Coordination with PG synthesis.** WthA’s strongest detected protein interaction was with the undecaprenyl phosphate recycling protein, UppP. Suggesting that WthA interaction network is even larger coordinating PG synthesis, turnover, and lipid carrier recycling.

Suppressor analyses in *A. baumannii* provided further insight into the WthA interaction network. Mutations in *mepM* or *mltD* rescued growth defects of the *wthA* mutant, suggesting that both enzymes become hyperactive in the absence of WthA. Increased MepM protein levels explain its elevated activity, whereas the mechanism controlling MltD may be either through direct negative regulation by WthA or indirect via coupling to MepM activity. These observations support a model in which WthA restrains multiple PG hydrolases to balance synthesis and degradation. This coupling is reminiscent of the classic observations that autolysis occurs when hydrolysis proceeds faster than synthesis, a principle that underlies β-lactam-induced lysis (28, 53–56).

Importantly, WthA’s impact on cell division appears distinct from LbcA-like activities. *P. aeruginosa* mutants lacking *lbcA* show no morphological abnormalities (51, 52), and in *A. baumannii*, deletion of *mepM* or *mltD* only partially restore growth and did not eliminate chaining or branching in Δ*wthA* cells. This separation of phenotypes indicates that WthA carries out additional functions— specifically, a role in cell division that is beyond its role in controlling PG hydrolyases (MepM, MltD).

Therefore, WthA may combine two levels of PG coordination that in other organisms are divided among multiple proteins reinforcing the idea that WthA is a multifunctional hub.

Homologs of WthA/LbcA are widely distributed across a α-, β-, and γ-Proteobacteria, as well as Bacteroidota and Chlorobiota (**Fig. 7B**), suggesting that these proteins represent a broadly conserved strategy for coordinating PG turnover. Notably, several of these homologs have been implicated in maintaining cell envelope integrity in *Brucella melitensis* (74), *Brucella abortus* (75), and *Rhizobium leguminosarum* (76), suggesting that WthA/LbcA-like proteins may perform related functions in other genera. However, it is unlikely that all homologs participate directly in cell division. In support of this notion, we found no evidence that *Pseudomonas* LbcA could influence the activity of its native amidase AmiB, the *P. aeruginosa* β-lactamase Oxa10, or the putative *A. baumannii* amidase, Oxa51 (**Fig 7D**). These findings underscore that WthA and LbcA differ in the range of PG turnover enzymes they regulate, reflecting functional diversification within this protein family (see **Fig. S7B-C**). How conserved each of these functions are across Gram-negative WthA/LbcA homologs will be interesting to explore.

Hub proteins that coordinate multiple PG turnover enzymes have been described in other bacteria. Among these, the roles proposed for *Acinetobacter* WthA most closely resemble those of *Caulobacter* DipM. DipM has been shown to function as a hub protein (69, 71), activate a cell division amidase (69–71), and interact with lytic transglycosylases (71). Similarly, *Pseudomonas* LbcA shows striking parallels to *Escherichia* NlpI as both employ TPR domains to direct proteolysis of endopeptidases through tail-specific proteases, CtpA in *Pseudomonas* (52, 77, 78) and Prc in *Escherichia* (79, 80). Together, these comparisons highlight a recurring theme that different species encode for multi-enzyme complexes that coordinate PG synthesis and hydrolysis to maintain envelope integrity.

It is also curious that LbcA and WthA differ in cellular localization, with LbcA anchored to the OM and WthA residing in the periplasm. All characterized amidase activators known to play major roles in cell division (EnvC, DipM, WthA) are soluble periplasmic proteins, whereas OM anchored activators (NlpD, ActS) exert more limited effects. This distinction may reflect the timing of key steps during cytokinesis in that periplasmic activators access nascent septal PG earlier, while OM-anchored factors act later as the OM invaginates. Consistent with this idea, all organisms possessing WthA/LbcA homologs but lacking EnvC encode soluble periplasmic variants (**Fig. 7B)**. Altogether, our findings define WthA as a previously unrecognized periplasmic hub protein that coordinates multiple PG turnover enzymes to promote balanced envelope remodeling during growth and division in *A. baumannii*.

## Materials and Methods

### Strains and growth conditions

All strains and plasmids used in this study are listed in **Dataset S1**. All cultures and agar plates were performed with Difco^TM^ Luria-Bertani Broth or Agar, Miller (BD). Where appropriate media was supplemented with glycerol (10 % volume/volume), kanamycin (30 μg/mL), tetracycline (5-10 μg/mL), hygromycin B (250 μg/mL for *A. baumannii* and 100 μg/mL for *E. coli*), carbenicillin (100 μg/mL), and anhydro-tetracycline (10 nM, aTc). Concentrations of isopropyl β-D-1-thiogalactopyranoside (IPTG) are indicated in the experiment.

*A. baumannii* 19606 deletion mutants were built by recombineering as described previously (81). The isogenic parent to 19606 recombineered strain, BWS399, was used as the 19606 “wild-type” for this study (11). TaKaRa ExTaq^®^ polymerase or Apex^TM^ Taq RED polymerase were used for PCRs <5 kb for cloning or screening experiments, respectively. PFU Turbo polymerase AD (Agilent) was used for PCRs >5 kb.

### Strain construction

An anhydro-tetracycline inducible mutant was built by the same recombineering methodology, but a hygromycin B resistant plasmid, pMMB67EHhph-RecAb, was used for induction of the RecAb system. Primers with the 125 nt of chromosomal homology were used to amplify from the plasmid pBaM3 which contained an outward facing tet-inducible promoter with an upstream and reverse orientation *tetR* regulator and kanamycin resistance cassette flanked by *frt* sites. The carbenicillin resistant pMMB67EH-Flp plasmid was used for removing *kan* cassettes from chromosomal inducible mutants. For the 19606 *tetR-wthA* strain, first the *frt-kan-frt-tetR-ptet* product (described above using pBaM3 template) was recombineered upstream of *wthA* on plates containing 10 nM aTc. Strains with inducible alleles were maintained with 10 nM aTc at all times. After the *kan* cassette was removed, a second *kan* cassette and ribosome binding site was recombineered downstream of *wthA* to give *lolB* a separate promoter and ribosome binding site using the 1334-cis-comp recombineering primers (pKD4 template). *tetR-wthA* was combined with *nlpD* and *tolA* deletions using the same methodology, but starting in 19606rec Δ*nlpD::frt* (pMMB67EHhph-RecAb) and 19606rec Δ*tolA::frt* (pMMB67EHhph-RecAb) strains, respectively.

### Plasmid construction

pMMB67EHhph-RecAb was built by amplifying the backbone of pMMB67EH-RecAb with XhoI-pMMBKn-for and SpeI-pMMBKn-rev. The *hph* resistance cassette was amplified using a lysate of a 5075 T101 mutant from the 5075 ordered library (45) as template and the primers SpeI-hph-for and XhoI-hph-rev. Inserts were digested with the restriction enzymes indicated by their primer names. Insert and vectors were ligated by T4 DNA Ligase (NEB) and transformed by electroporation into DH5α competent cells.

pBAD18kn-express was built by amplifying the *frt* flanked *kan* cassette from pKD4 with ClaI-kan-for and ClaI-kan-rev. pBAD18 vector and insert were digested with the restriction enzymes indicated by their primer names. Vector was treated with alkaline phosphatase. Insert and vectors were ligated by T4 DNA Ligase (NEB) and transformed by electroporation into DH5α competent cells.

pBaM1 was built by amplifying the backbone of pMMB67EHkan with primers PacI-pMMB-swap-for and PacI-pMMB-swap-rev. The insert *frt-kan-frt-araC*-pBAD promoter was amplified from pBAD18kn-express with primers PacI-pBAD-fr and PacI-pBAD-rv. Inserts were digested with the restriction enzymes indicated by their primer names. Insert and vectors were ligated by T4 DNA Ligase (NEB) and transformed by electroporation into DH5α competent cells.

pBaM3 was built by amplifying the *tetR* regulator and *tetR/tetA* promoter from p-dcas9-bacteria (82) with primers ClaI-ptet-for and EcoRI-ptet-rev. pBaM1 vector and insert were digested with the restriction enzymes indicated by primer names. Vector was treated with alkaline phosphatase. Insert and vectors were ligated by T4 DNA Ligase (NEB) and transformed by electroporation into DH5α competent cells.

*Pseudomonas aeruginosa* PA01 alleles of *lbcA* and *amiB*, *lbcA* and *oxa51*, *nlpD* and *amiB*, and *nlpD* and *oxa51* were codon optimized for expression in *E. coli* (GenScript). Gene synthesized products were designed to have an EcoRI cut site, AGGAGA ribosome binding site, putative activator (*lbcA* or *nlpD*), KpnI cut site, AGGAGA ribosome binding site, putative amidase (*amiB* or *oxa51*), and XbaI cut site. Gene synthesis was performed by GenScript in the pUC57mini plasmid.

Complementation and overexpression plasmids were built by digesting pMMB67EHtet with appropriate restriction enzymes (NEB) and treating digested plasmid with Antarctic Phosphatase (NEB). Inserts were digested with the restriction enzymes indicated by their primer names. Insert and vectors were ligated by T4 DNA Ligase (NEB) and transformed by electroporation into DH5α competent cells. Plasmids were screened by colony PCR using primers ptac-seq and rrnB-T1-term.

### Microscopy

Overnight cultures were diluted into fresh 5 mL cultures of appropriate media at OD_600_ ∼0.05. Strains were grown to OD_600_ ∼0.5-0.7 and 1 μl was spotted on VWR No. 1.5, 50X22mm rectangular micro cover glass (48393–195). Difco Minimal Agar Davis (254410) pads were placed on top of samples and imaged camera at 100x magnification on an inverted Nikon Eclipse TI2-E equipped with an Orca-Fusion Gen-III sCMOS camera and CFI60 Plan Apochromat Lambda Phase Contrast DM 100x Oil Immersion Objective Lens. NIS Elements software was used to acquire Z-stack images with a range of 1um, 7 steps per capture, 0.2 um per step. Three plains of each Z-stack were averaged with a Z-projection using ImageJ. Z-projections assisted in viewing of the highly kinked chains of *wthA* mutants which often extended slightly up and down along the Z-plane even with the flattening effects of agar pads. For consistency, all microscopy was performed and processed in the same manner.

Cell membranes and late-stage division sites were visualized by staining with FM™ 4-64 Dye (N-(3-triethylammoniumpropyl)-4-(6-(4-(diethylamino) phenyl) hexatrienyl) pyridinium dibromide, Invitrogen™). A 100 μg aliquot of dye was dissolved in 50 μl of DMSO, vortexed, and stored at room temperature in the dark during the experiment. 500 μl of cell culture were mixed with 5 μl of fresh dye, vortexed briefly, and incubated for 15 min at room temperature in the dark. Cells were pelleted at 5,000 x *g* for 3-5 min and washed once with 500 μl of 1x PBS. Cells were pelleted again and resuspended in 100 μl of 1x PBS. Cells were imaged for both phase contrast and FM™ 4-64 fluorescence as described above.

Samples were prepared for transmission electron microscopy by diluting overnight cultures into fresh 5 mL of appropriate media at OD_600_ ∼0.05. Strains were grown to OD_600_ ∼0.7 and 5 mL of culture was pelleted at 5000 x g for 7 min. Cells were washed once with 1x PBS (Fisher Bioreagents) and pelleted again. Pellets were resuspended in a fixative mixture containing 2 % of EM grade glutaraldehyde (Electron Microscopy Sciences) and 0.1 M trihydrate sodium cacodyate (Electron Microscopy Sciences) pH 7.25 with HCl. Samples were submitted to Georgia Electron Microscopy for transmission electron microscopy.

### Growth curves

Overnight cultures were diluted into fresh 5 mL of appropriate media at a starting OD_600_ of 0.05. Cultures were incubated at 37 °C until stationary phase was apparent, when a dip in OD_600_ indicated that lysis may occur the cultures were continued for 1-2 hours to see if this drop continued. At each timepoint, 200 μl of culture was transferred to wells in a 96-well flat-bottom polystyrene costar^®^ microtiter plate (Corning) to measure OD_600_ on a Synergy H1 Hybrid reader with Gen5 software (BioTek^®^).

Overnight cultures for depletion of *wthA* contained 10 nM aTc. To assist in depletion of the tet-inducible *wthA* allele, cultures were diluted as described above into fresh 5 mL of LB with appropriate supplements. At the 2-hour time point, 50 μl of each culture was diluted again into a fresh 5 mL of the same media to perform a single back-dilution.

### Sensitivity to cold storage assessment

Strains were struck on two separate LB plates. After incubation at 37 °C one plate of each was stored at room temperature and one plate at 4 °C for 7-8 days. Single colonies from each storage condition were restruck on LB plates and incubated at 37 °C. The experiment was performed in triplicate for all strains.

### Minimum inhibitory concentration (MIC) determination

For *A. baumannii* strains with altered WthA expression, overnight cultures were grown in biological triplicate for each strain in 5 mL of LB with appropriate supplements. To assist in depletion of tet inducible strains, overnight cultures were grown with no aTc. Cotton-tipped applicator sticks were used to spread a lawn of each overnight on LB agar plates and the indicated E-test strip (Liofilchem®) was added to each lawn. Plates were incubated at 37 °C for ∼16 hours and the MIC determined by where the lawn bisects the E-test strip.

For *E. coli* strains overexpressing Oxa51 or a TEM-1 β-lactamase, MICs were assessed by growth in 100 μl cultures in LB broth in each well of 96-well flat-bottom polystyrene costar^®^ microtiter plates (Corning). Briefly, overnight cultures were grown in biological triplicate for each strain in 5 mL of LB with appropriate antibiotics for plasmid maintenance. Overnights were diluted 1 in 1,000 into fresh LB broth and dispensed into wells of a microtiter plate. Antibiotics were added in an additional 100 μl of media to the first column of each plate. Antibiotics were serially diluted down the plate by moving the additional 100 μl of volume to the next column, leaving the final column with no drug added. Plates were sealed with AeraSeal^TM^ film (EXCEL Scientific, inc.) and incubated at 37 °C for ∼16 hours. OD_600_ was measured on a Synergy H1 Hybrid reader with Gen5 software (BioTek®). MIC determinations were calculated as the concentration at which 90% of growth was inhibited relative to the no drug controls. Error indicates the standard deviation of the biological triplicates.

### Suppressor isolation

Overnight cultures of 19606 Δ*wthA* were grown in LB. 50 μl of each overnight was diluted in fresh 5 mL of LB with 5% glycerol and incubated at 37 °C for 24 hours. 50 μl of each 5% glycerol culture was diluted in fresh 5mL of LB with 10 % glycerol and incubated at 37 °C for 24 hours. 1 mL of cells was pelleted at max speed for 1 min for cultures with high optical density after growth in LB with 10% glycerol. The supernatant was removed, and pellets were whole genome sequenced as described above at SeqCenter, LLC.

### Immunoblots and pull-down assays

To assess protein levels of MepM-His, cells were cultured, lysates were prepared, and 10 μg of protein was run on a NuPAGE® 10% Bis-Tris gel (Invitrogen) as described (60). Proteins were transferred to 0.45 μm nitrocellulose (Amersham™ Protran™ Premium) as described (60) and probed with THE™ His Tag Antibody, mAb, Mouse (Genscript) as the primary antibody. Cy5™ goat anti-mouse IgG (Invitrogen) was used as the secondary and blots were imaged for on a BioRad ChemiDoc™ MP imaging system. Proteins for HA and His pulldowns described below were loaded at the indicated volumes and blots prepared as described below. the™ HA-tag antibody mouse (Genscript) and MonoRab™ Anti-His Tag antibody rabbit (Genscript) were used as primary antibodies. Cy5™ goat anti-rabbit IgG (Invitrogen) and Cy3™ goat anti-mouse IgG (Invitrogen) were used as secondary antibodies and image as described above.

For pull down assays, overnight cultures were grown in 5 mL of LB broth with 5 μg/mL of tetracycline. Overnights were diluted to a starting OD_600_ of ∼0.05 in 50 mL of LB broth with 5 μg/mL of tetracycline and 25 μM IPTG and grown at 37 °C until the cultures reached an OD_600_ ∼1.0. 50 mL of cells were pelleted at 5,000 x *g* for 20min at 4 °C and resuspended in 5 mL of 1x PBS. The sample was split to two tubes (one for His and one for HA pull downs) and centrifuged again. The supernatants were removed and ∼100 mg of pellets were frozen at-80 °C. Pellets were thawed and resuspended in 1 mL of lysis buffer containing 1 x BugBuster® Protein Extraction Reagent (Millipore), 0.5 mg/mL rlysozyme (Millipore), 2 mM CaCl_2_, 0.1% triton X-100, 1 x Benzonase® Nuclease (Millipore), and cOmplete™ Mini EDTA-free Protease Inhibitor Cocktail (1 tablet per 10mL of lysis buffer). Lysates were rotated at room temperature for 3 hours. Remaining cell debris was removed by centrifuging at 17,000 x *g* for 10 min and the supernatant transferred to a fresh tube.

HA-pulldowns were performed following the guidelines of the Pierce Magnetic HA-TagIP/Co-IP Kit (ThermoScientific) with the following adjustments. 25 μl of beads were mixed with 975 μl of lysate described above and incubated on rotator for 2 hours at room temperature. 0.2 % Triton X-100 was added to the provided Lysis/Wash Buffer and used for three washes followed by a fourth wash with 300 μl of sterile ddH_2_O. Proteins were eluted with 50 μl of 5 mM HA peptide (Genscript) and incubated on rotator for 30 min at room temperature.

His-pulldowns were performed following the guidelines provided for HisPur™ Ni-NTA Resin (Thermo Scientific™) with the following adjustments. 987 μl of lysate were mixed with 70 μl of 1 M HEPES (final concentration 50 mM), 140 μl of 5 mM NaCl (final concentration 500 mM), 175 μl of 80% glycerol (final concentration 10%), and 28 μl of 1 M imidazole (final concentration 20 mM). The lysate mixture was rotated for 1 min to ensure it was mixed with 100 μl bed volume of prepared resin in a capped TALON® 2 ml Disposable Gravity Column (TaKaRa). Lysate and resin were rocked for 2 hours at room temperature. Resin was washed with a total of 10x the bed volume of wash buffer containing 50 mM HEPES, 500 mM NaCl, 20 mM imidazole, 10% glycerol, and 0.2% triton X-100. The resin was eluted stepwise with 500 μl each of 100-, 200-, and 500 mM imidazole in 50 mM HEPES, 500 mM NaCl, and 0.2% triton X-100. Elutions were combined and precipitated with a final concentration of 33% trichloroacetic acid (TCA) on ice at 4 °C for ∼16 hours. TCA precipitates were centrifuged at 17,000 x *g* for 30 min at 4 °C and washed twice with 1 mL of ice cold acetone. Washes were centrifuged at 17,000 x *g* for 15 min. After the second wash, precipitates were air dried for 2 min and resuspended in 50 μl of 1 x NuPAGE sample buffer (Invitrogen) with 5 % 2-mercaptoethanol.

For mass spectrometry identification of proteins interacting with HA-WthA, a 30 mL culture was grown as described above to OD_600_ ∼1.0. 25 mL of culture was centrifuged at 5,000 x *g* for 7 min at 4 °C. Cells were washed twice with 5 mL of 1 x PBS and re-pelleted. The cell pellets were resuspended in 920 μl of 1x PBS and mixed with 80 μl of freshly prepared 25 mM EGS (ethylene glycol bis(succinimidyl succinate), Thermo Scientific™) dissolved in DMSO. Cells were incubated with EGS cross-linker for 30 min at room temperature and then quenched with 5 μl of 1 M Tris, pH 7.5 at room temperature for 15 min. Cross-linked cells were spun down at 17,000 x *g* for 1 min and washed once with 1 mL of 50 mM Tris, pH 7.5. Cross-linked pellets were frozen at-80 °C and lysates prepared as described above. Proteins were pulled down using the Pierce Magnetic HA-TagIP/Co-IP Kit (ThermoScientific) as described above except proteins were eluted using the kit’s elution and neutralization buffers following guideline of the manual. Elutants were TCA precipitated as described above, but precipitates were not resuspended and were frozen at-80 °C. Precipitates were submitted to The Georgia Institute of Technology Systems Mass Spectrometry Core for shotgun proteomics with trypsin digest. Peak abundances for each protein identified in the shotgun proteomics was compared to abundance detected in two negative controls, a strain with an empty plasmid and a strain expressing the cytoplasmic protein ElsL-HA. Hits were filtered for having a >5 fold increased peak abundance compared to both controls, a p-value and q-value <0.01, >1 unique peptide detected, >10% coverage of the protein sequence, and correct periplasmic localization for biologically relevant interactions.

### Methods for demonstrating PG amidase activity

#### PG Amidase Release Assay

The strain *E. coli* W3110 Δ*oppBCDF::frt* Δ*cadB::frt* was used for assays for release of PG amidase products (60, 61). The strain (“AmpG-only” strain) is missing both transporters for recycling of amidase products (freed peptides) but retains the AmpG transporter that allows for recycling the products of endopeptidases and lytic transglycosylases. Cultures of smaller volumes (2-5 ml) for radioactive assays were grown in glass tubes, whereas 30 mL cultures were grown in 125 mL sterile disposable PETG plain bottom flasks with vented closures (Thermo Scientific). As described previously (60), [^3^H]-Dap (American Radiolabeled Chemicals Inc., (DL+meso)-2,6-diaminopimelic acid [2,6-^3^H], 60 Ci/mmol, 1 mCi/mL stock) was used to label the PG peptide in 2 mL overnight cultures of LB with 500 nCi/mL of [^3^H]-Dap and 5 μg/mL of tetracycline. 50 μl of overnight cultures was diluted into fresh 5 mL of LB with 500 nCi/mL of [^3^H]-Dap, 5 μg/mL of tetracycline, and the indicated concentrations of IPTG. Cultures were grown at 37 °C for 3-4 hours to OD_600_ ∼1.0. The 5 mL cultures were transferred to 15 mL conical tubes and centrifuged at 5,000 x *g* for 5 min. Supernatants were discarded and cell pellets were resuspended in fresh 5 mL of LB with tetracycline and IPTG. The 5 mL resuspensions were diluted into total volume of 30 mL of fresh, pre-warmed LB containing tetracycline and IPTG, and a final concentration of 30 μg/mL of cold mDap (2,6-diaminopimelic acid, Sigma). A 0-hour time point was taken, and strains were grown at 37 °C for a total of 3 hours to assess release of amidase products. During growth, cultures were removed every hour and processed as follows: (i) the OD_600_ was read in cuvettes, and (ii) 500 μl of each sample was transferred to a 1.5 mL centrifuge tube and centrifuged at 17,000 x *g* for 2 min. Supernatants, containing [^3^H]-Dap that had been released from the cell, were carefully pipetted off and pellets, containing cellular [^3^H]-Dap that had been retained and/or recycled, were resuspended in 50 μl of PBS. 10 μl of the cellular resuspension was diluted into 5 mL of ScintiVerse™ BD cocktail (Fisher) in scintillation vials and vortexed. Liquid scintillation was counted by a Tri-Carb^®^ 4910 TR liquid scintillation counter (PerkinElmer) with QuantaSmart™ program. The percent of tritium products released was calculated by percent of tritium lost comparing the 0-and 3-hour time-points.

#### Lysis controls for amidase assays

To assess if lysis occurred during the amidase product release assays described above, 20 μl of sample from each time point during the chase was diluted into 180 μl of LB in the first row of a microtiter plate. A 1 in 10 dilution series, total volume of 200 μl, was performed down the row of the plate. 4 μl of each well was spotted on LB agar plates (1 in 250 plating dilution). Once spots dried, plates were incubated at 37 °C for ∼16 hours and cfu’s were counted at the lowest dilution that had ∼10x cfu’s in the preceding dilution. Cfu/mL were calculated. In addition, only at the 3-hour final timepoint of the PG amidase release assay, a portion of the cell supernatant (400 μl) was stored for assessment of DNA or ATP released. In technical duplicates of biological triplicate, 100 μl each of the supernatants was assessed for DNA or ATP release to the supernatant following the respective manuals for LIVE/DEAD® BacLight™ Bacterial Viability Kit (Invitrogen) propidium iodide fluorescence in microtiter plates and BacTiter-Glo™ Microbial Cell Viability kit (Promega) luciferase luminescence in microtiter plates. Assays were read in Corning™ 96-Well Solid Black Polystyrene Microplates on a Synergy H1 Hybrid reader with Gen5 software (BioTek^®^).

#### Amidase Uptake Assays

To determine whether the products released in the PG amidase release assays were indeed freed peptides, we adapted a previously published MppA/OppBCDF scavenging assay (60). This transport system has strict specificity for freed PG peptides. The strain background used for the Amidase Uptake Assay, that we called the “Opp-only” strain, included several genetic changes. (i) the strain contains a plasmid-expressing OppBCDF to specifically import freed PG peptides (amidase products). (ii) The gene encoding AmiD, which functions as a periplasmic PG recycling amidase that can cleave PG peptides attached to anhydro-MurNAc (endopeptidase and lytic transglycosylase products), has been deleted. Without AmiD we have shown that anhydro-MurNAc linked PG peptides cannot be redirected to MppA/OppBCDF (60). (iii) The strain also lacks *ampG* which encodes the permease for importing anhydro-MurNAc linked PG peptides. (iv) Finally, the Opp-only strain lacks *ampD* which encodes the cytoplasmic PG recycling amidase. We have previously observed that in the absence of AmpD, the accumulation of anhydro-MurNAc linked PG peptides in the cytoplasm leads to a feedback inhibition preventing further uptake (60). Therefore, even if anhydro-MurNAc linked PG peptides somehow were imported in our Opp only strain they would block further uptake of these products.

Spent media were collected at the end of the release assays described above, filter-sterilized, refreshed with 0.2% glucose and 50mM NH_4_Cl. Uptake assays were performed as described previously (Scavenging assay protocol in ref 5) except the Opp-only strain overnights were grown in LB broth and the spent media came from refreshed LB as described above.

### Bioinformatic analyses

A summary of the homolog search results is in in **Data Set S1**. All homologs were identified by blasting known genes from *E. coli* (for EnvC/NlpD/ActS), *A. baumannii* (for WthA), or *Caulobacter* (for DipM). Significant hits were all verified by blasting the hit back against these genomes to identify the gene it was most similar to, and by searching for literature that may have described an alternative function for the hit with PaperBlast (83). SignalP-5.0 (84) was used to check localization of each allele. A phylogenic tree was generated using PhyloT v2 and the tree was visualized and annotated with iTOL (85).

## Data availability

All data related to this paper are within the text, supplemental, or may be requested from the authors.

## Supporting information

Data Set S1

## Acknowledgments

We gratefully acknowledge funding from the National Institutes of Health, grants AI176776, AI138576, and AI150098 (to M.S.T.); funding from the Army Research Office (ARO, http://www.arl.army.mil/) grant W911NF2010195 (to M.S.T.); and grant F32 GM137554 (to B.W.S.).

## Supplemental Figures and Tables

**Fig S1.**
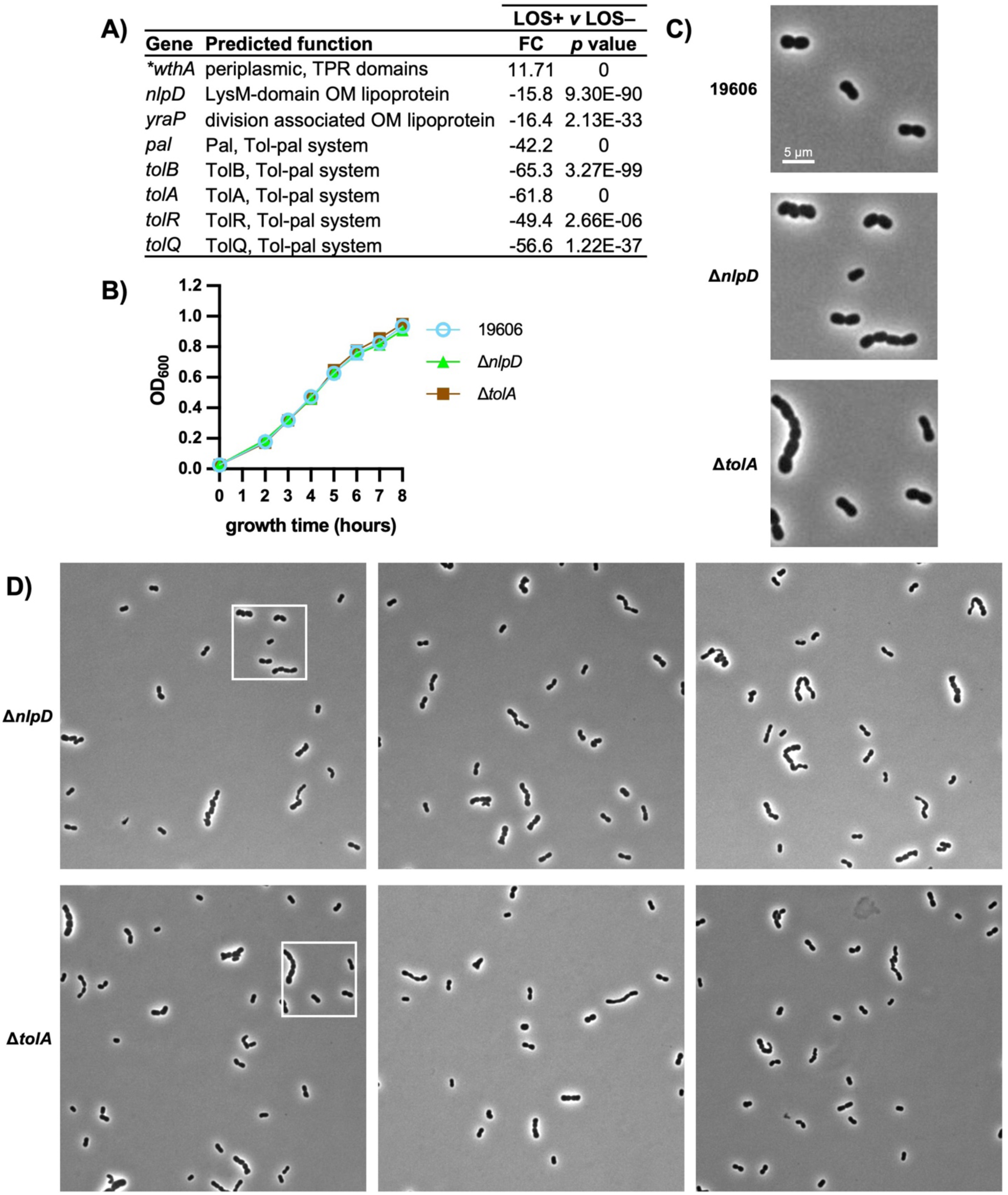
Phenotypes of *A. baumannii nlpD* and *tolA* mutants. (A) Select hits from our previous Tn-seq screen comparing genes critical for function in the presence and absence of LOS. NlpD, Tol-Pal, and YraP become critical for fitness in an LOS-deficient strain. *wthA* is easy to disrupt in LOS-deficient cells, perhaps because it does not function in the absence of LOS. FC indicates fold changes. (B) Biological triplicate growth curves comparing growth of *A. baumannii* 19606 wild-type to growth of *nlpD* and *tolA* mutants. Error bars are not shown when smaller than the size of the symbol. (C) Phase contrast microscopy of *nlpD* and *tolA* mutants, representative of biological triplicate. (D) Full-sized, additional fields of view for microscopy of *nlpD* and *tolA* mutants. First image shows a box for the cropped area shown in panel C.

**Figure S2.**
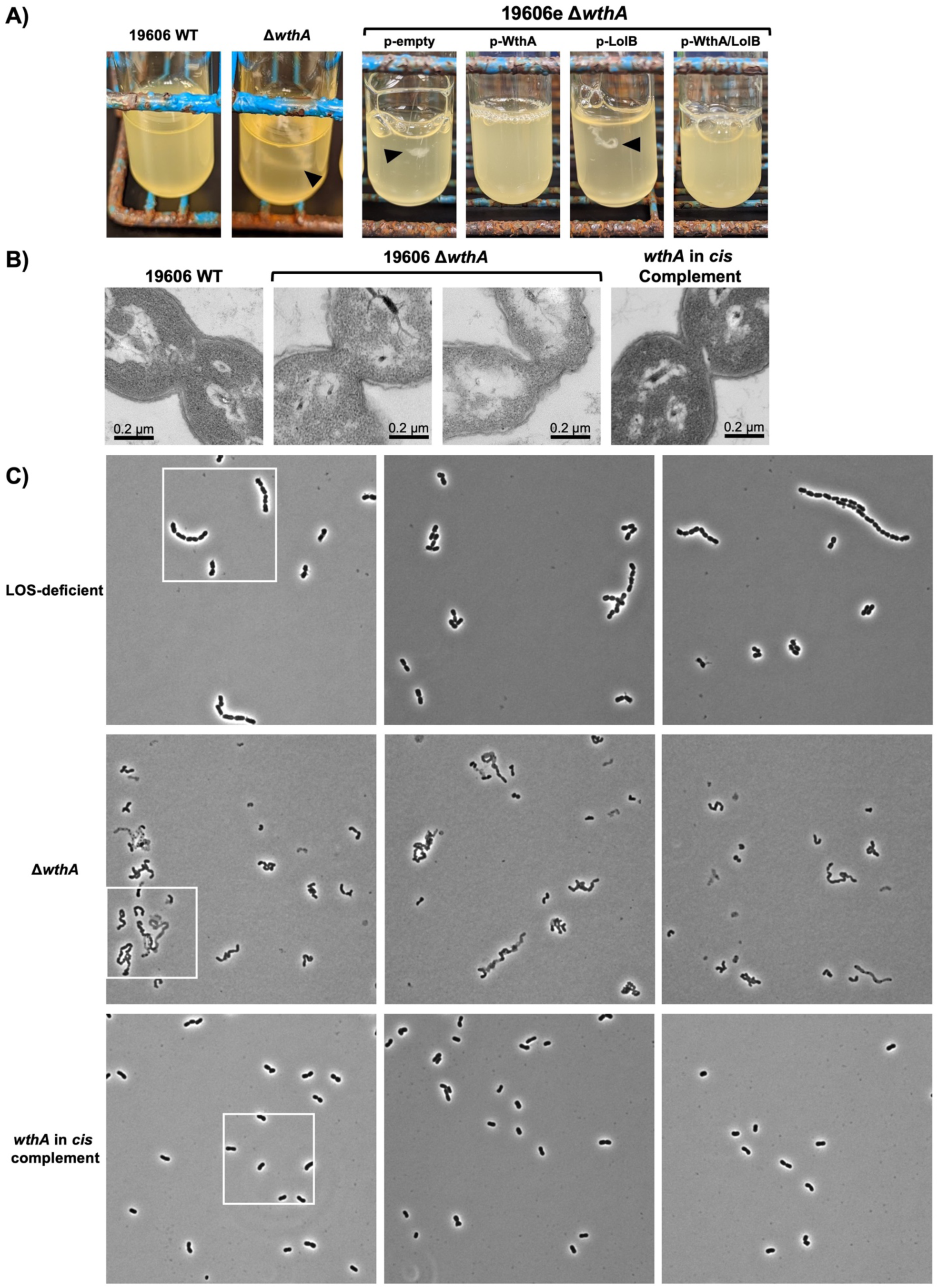
Cellular debris and microscopy of *wthA* mutants grown in LB. (A) Clumped debris material is indicated with black arrow-heads and is accompanied by low density of overnight cultures. All images are representative of growth trends observed in at least biological triplicate. 19606e, electrocompetent, background is described in **Fig. S3**. (B) Transmission electron microscopy focusing on division sites of *wthA* mutants. (C) Full-sized, additional fields for microscopy of LOS-deficient and *wthA* mutants. First image shows a box for the cropped area shown in **Fig. 2A**.

**Figure S3.**
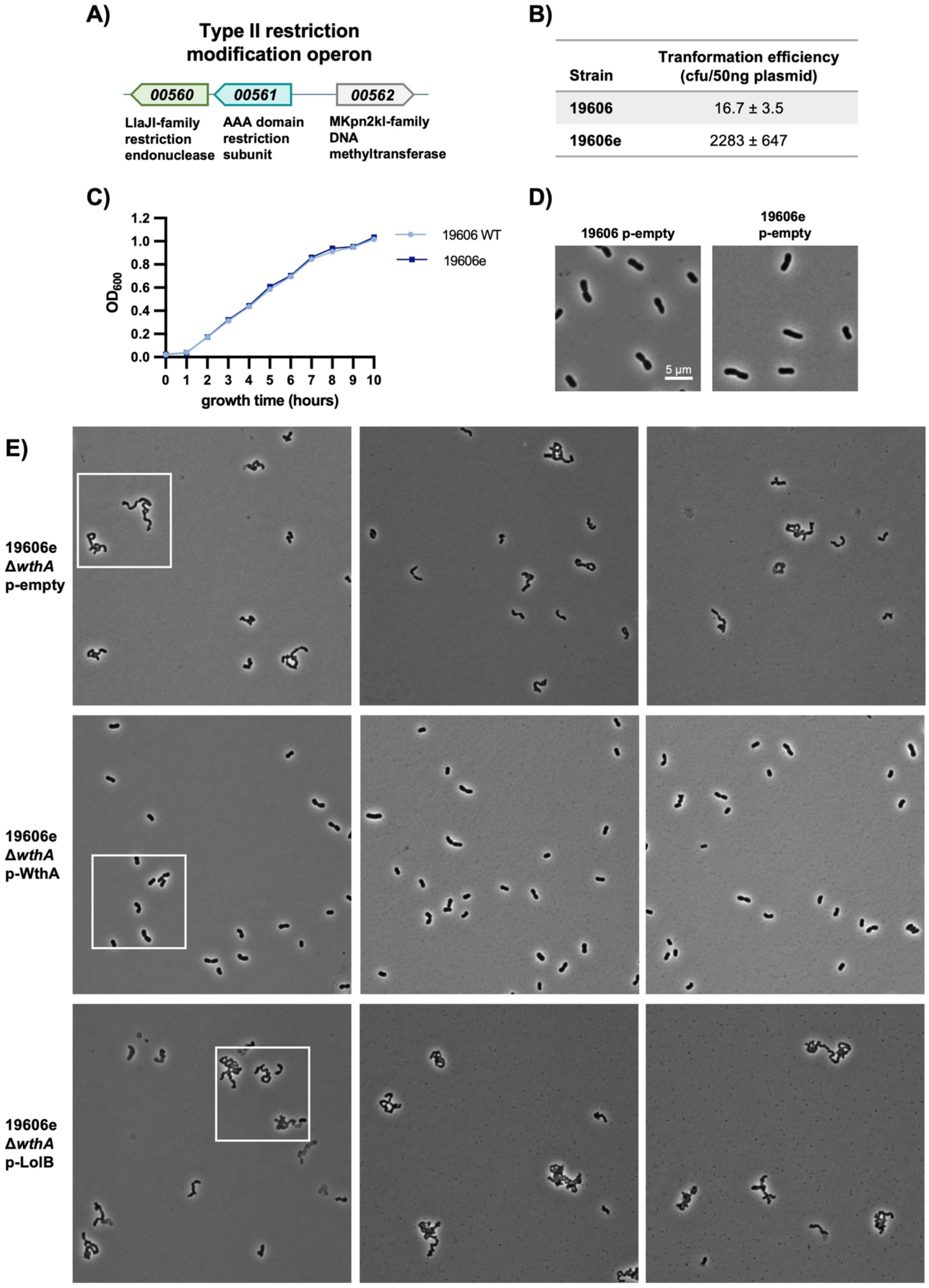
Generation of an electrocompetent strain of *A. baumannii* 19606 to test *wthA* complementation. (A) Operon structure of the HMPREF0010_00560-HMPREF0010_00562 restriction/modification system. (B) Transformation efficiency of 19606 and the electrocompetent 19606e (Δ00560-00561) strains, average and standard deviation of biological triplicates. (C) Growth curves of 19606 and 19606e strains in biological triplicate. Error bars are not shown when smaller than the size of the symbol. (D) Phase contrast microscopy of 19606 and 19606e strains, representative of biological triplicates. (E) Full-sized, additional fields for microscopy test of *wthA* complementation. First image shows a box for the cropped area shown in **Fig. 3B**.

**Figure S4.**
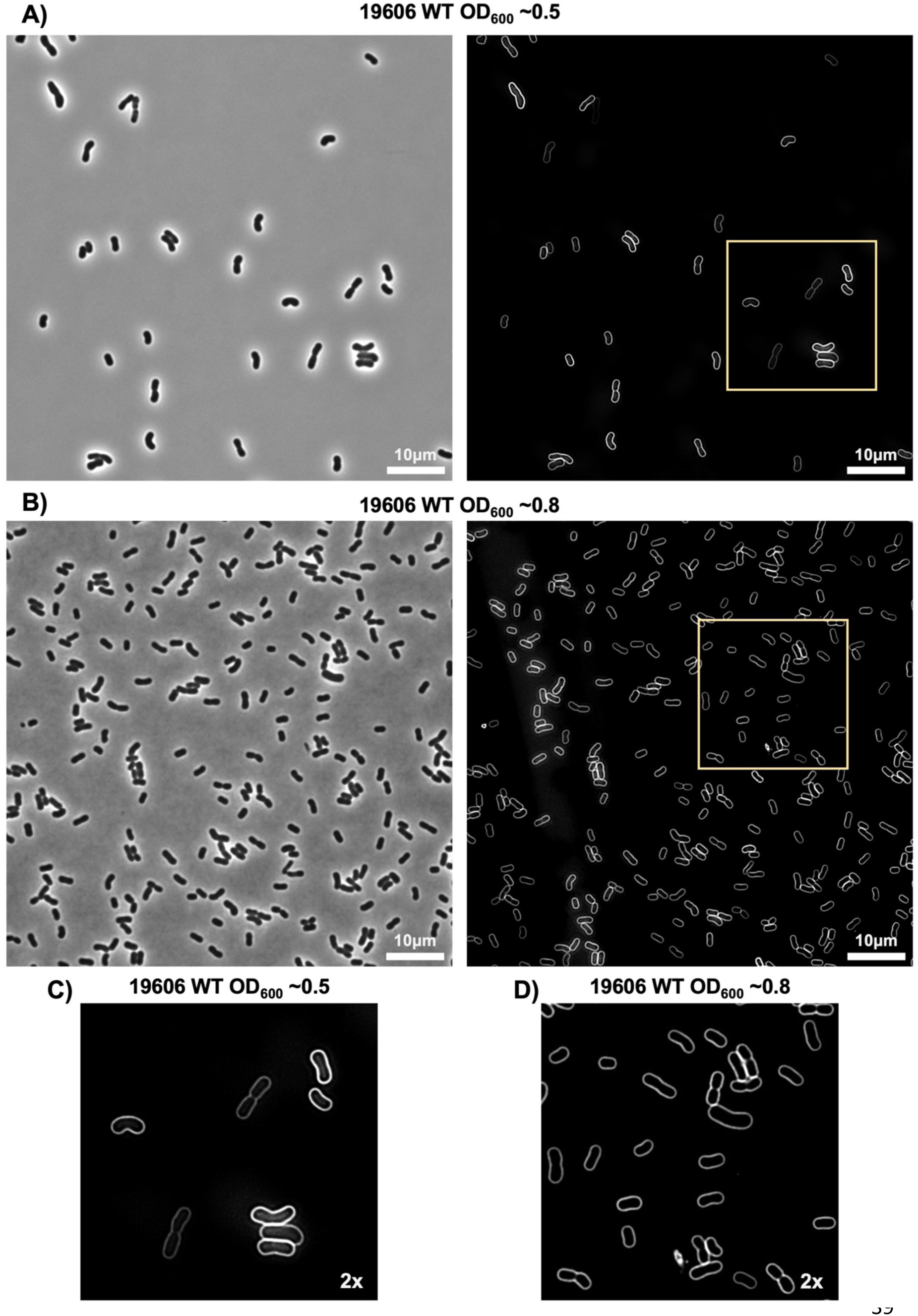
***A. baumannii* ATCC 19606 membrane staining.** Phase contrast imaging and imaging of membrane staining with FM 4-64 of the 19606 WT parent strain grown to mid logarithmic (A,C OD_600_ ∼0.5) and late logarithmic (B,D OD_600_ ∼0.8) stage. (A-B) Complete fields of view with phase contrast on left and FM-4-64 fluorescence on right. (C-D) 2x zoomed in regions (yellow box on full image) of the images above.

**Figure S5.**
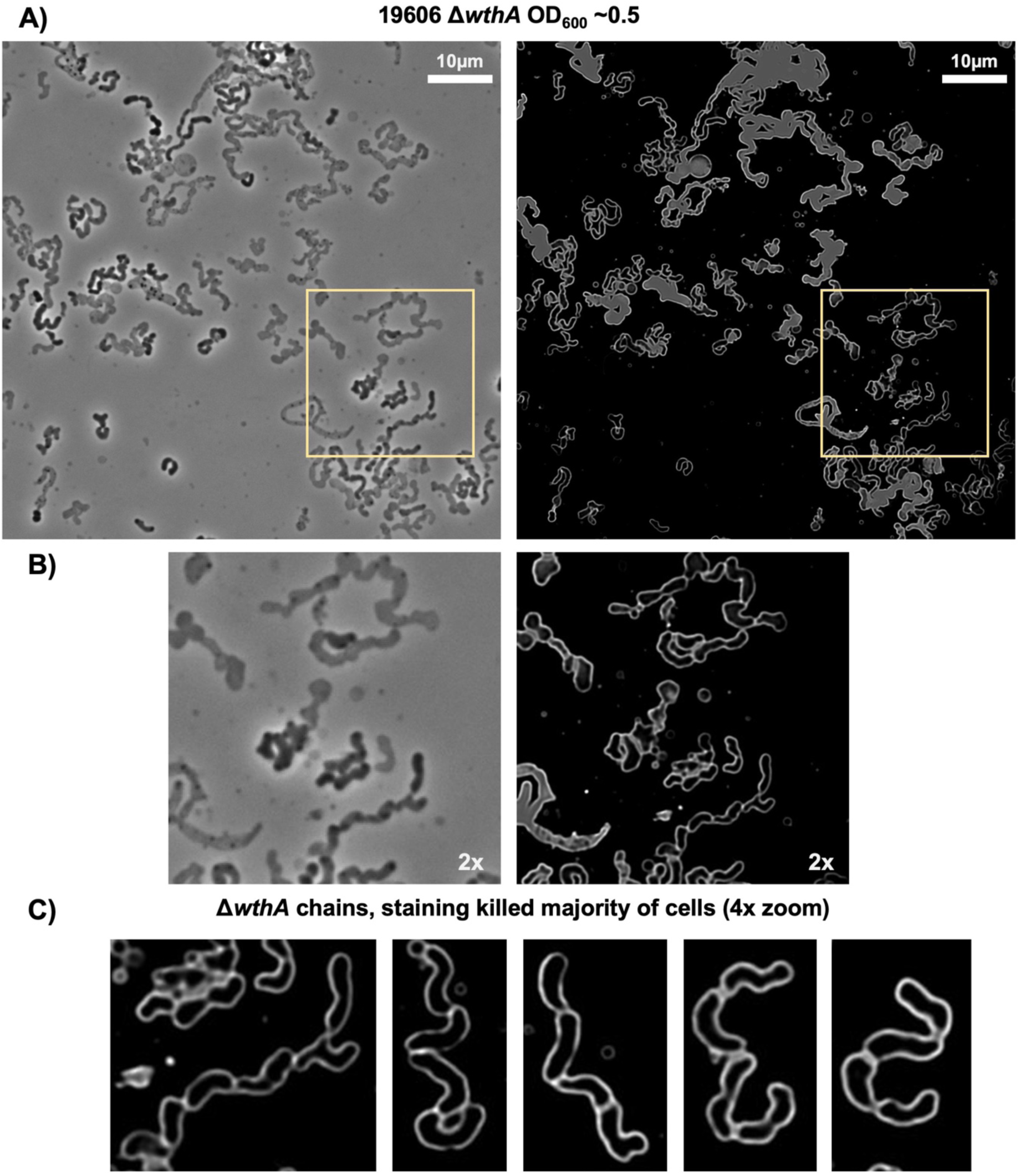
Morphological defects in the absence of WthA, mid logarithmic stage of growth. Membrane staining with FM 4-64 of 19606 Δ*wthA* cells during logarithmic stage of growth (OD_600_ ∼0.5). (A) Complete fields of view with phase contrast on left and FM-4-64 fluorescence on right. (B) 2x zoomed in regions (yellow box on full image) of the images above. (C) Additional 4x zoomed in examples of cells with chaining. All microscopy are representative of at least biological triplicates.

**Figure S6.**
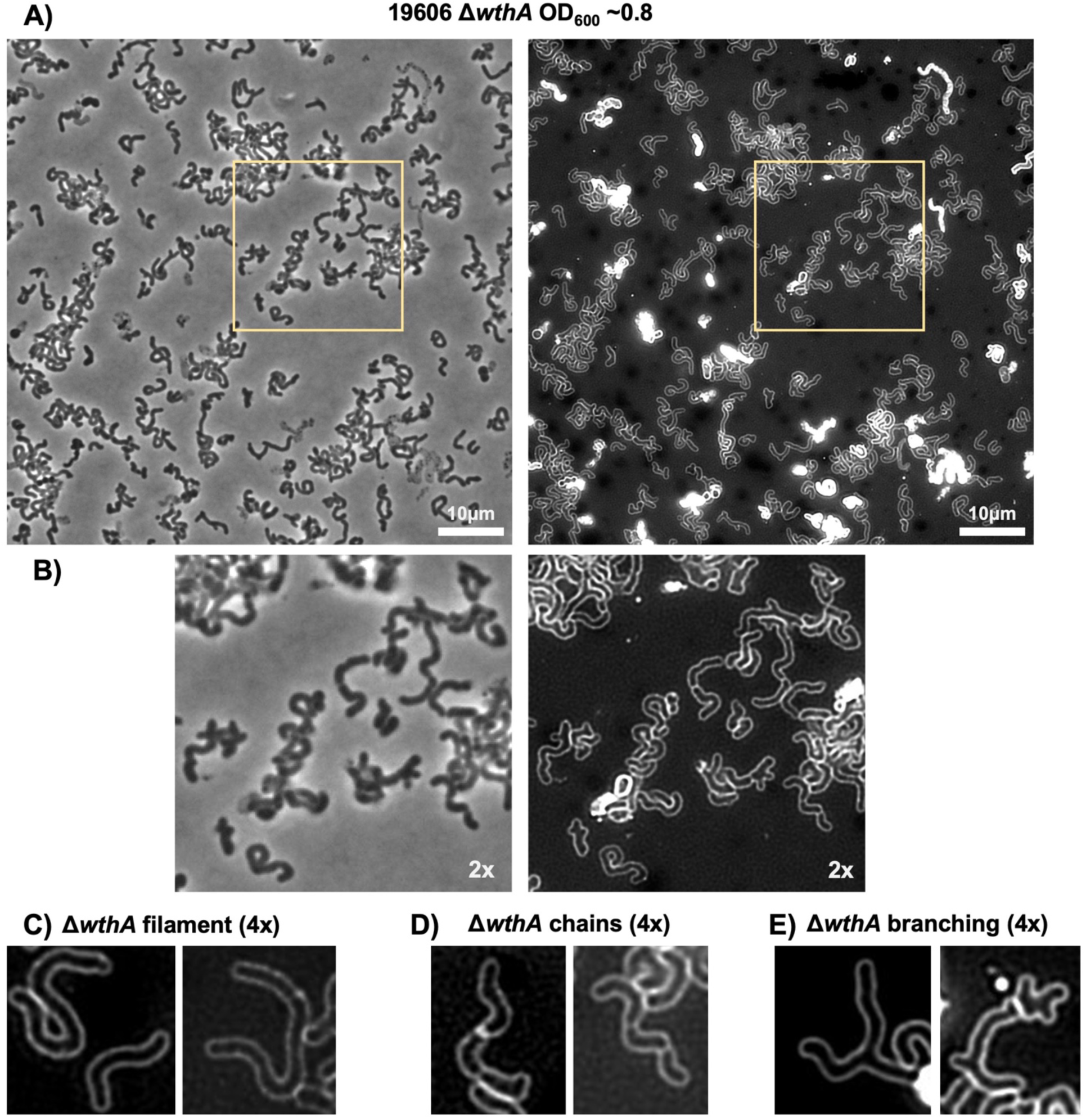
Morphological defects in the absence of WthA, late logarithmic stage of growth. Membrane staining with FM 4-64 of 19606 Δ*wthA* cells during late logarithmic stage of growth (OD_600_ ∼0.8). (A) Complete fields of view with phase contrast on left and FM-4-64 fluorescence on right. (B) 2x zoomed in regions (yellow box on full image) of the images above. (C-E) Additional 4x zoomed in examples of cells with 3 major morphological defects observed in the 19606 Δ*wthA* strain: filamenting (C), chaining (D), and branching (E). All microscopy are representative of at least biological triplicates.

**Figure S7.**
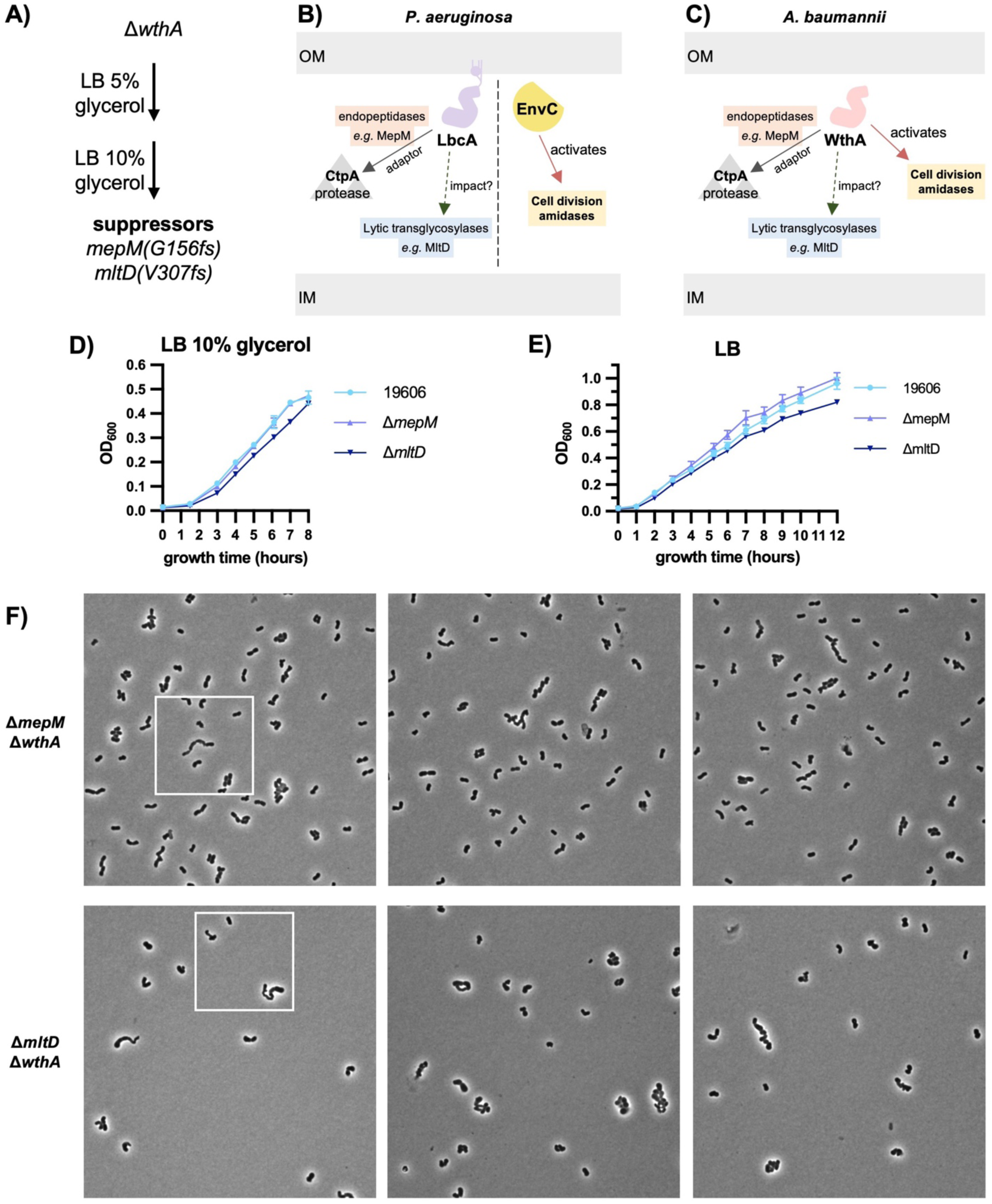
Effects of disrupting MepM and MltD in *A. baumannii*. (A) Strategy of 19606 Δ*wthA* suppressor selections with increasing osmolarity which resulted in two populations with the indicated mutations. (B) Summary of PG turnover enzymes impacted by LbcA (homolog of WthA in *Pseudomonas*) and EnvC. (C) Possible impact of WthA on PG turnover enzymes (D-E) Growth curves in LB + 10% glycerol (D) and LB (E) of *mepM* and *mltD* mutants. Data is from biological triplicate and error bars are not shown when smaller than the size of the symbol. (F) Full-sized, additional fields for microscopy of Δ*wthA* suppressors. First image shows a box for the cropped area shown in **Fig. 4F**.

**Figure S8.**
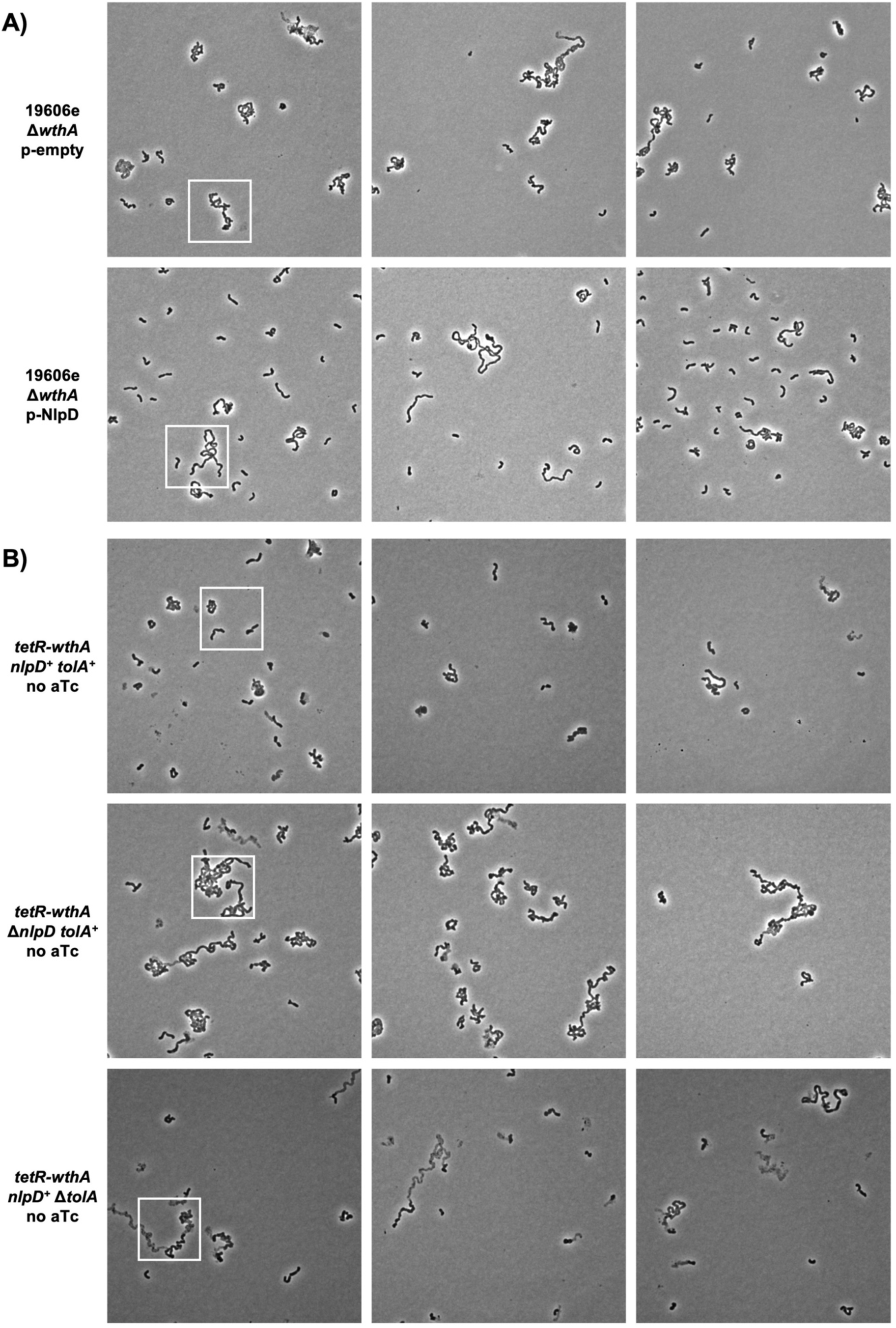
Additional microscopy of cells with alterations to WthA and NlpD. (A) Full-sized, additional fields for microscopy of 19606e Δ*wthA* with empty plasmid or plasmid overexpression of NlpD. First image shows a box for the cropped area shown in **Fig. 5B**. (B) Full-sized, additional fields for microscopy of *wthA* depletion in the presence or absence of NlpD and TolA. First image shows a box for the cropped area shown in **Fig. 5D**.

**Figure S9.**
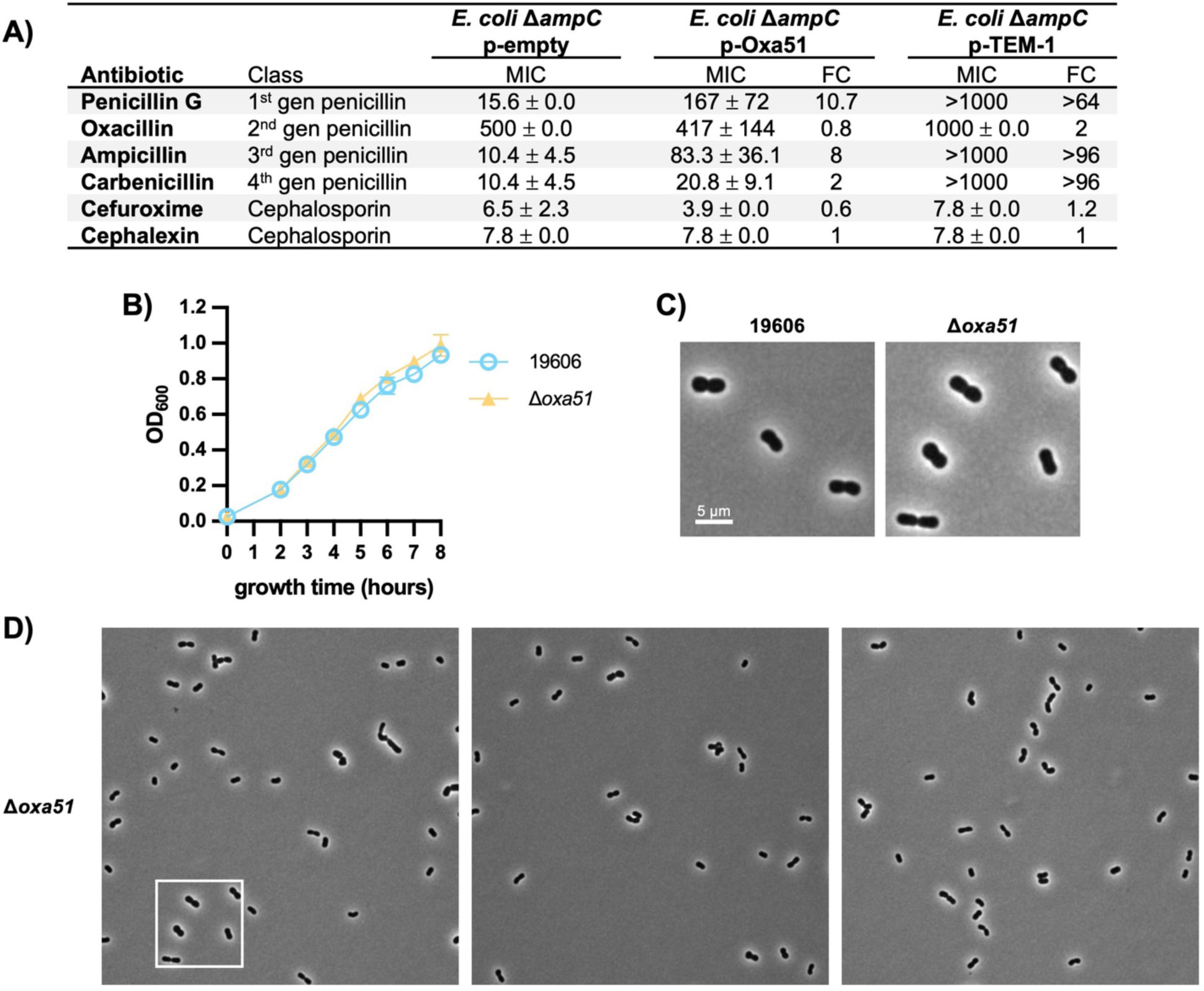
Loss of Oxa51 has no major impact on cell division. (A) Minimal inhibitory concentration of the indicated β-lactams determined in liquid cultures for an *E. coli* strain with the indicated plasmids. Expression in an *E. coli* strain was used because it had fewer chromosomally encoded β-lactamases. (B) Growth curves comparing growth of *A. baumannii* 19606 wild-type to growth of an *oxa51* mutant. Data is from biological triplicate and error bars are not shown when smaller than the size of the symbol. (C) Representative phase contrast microscopy of an *oxa51* mutant, representative of biological triplicate. (D) Full-sized, additional fields for microscopy of the *oxa51* mutant. First image shows a box for the cropped area shown in (C).

**Figure S10.**
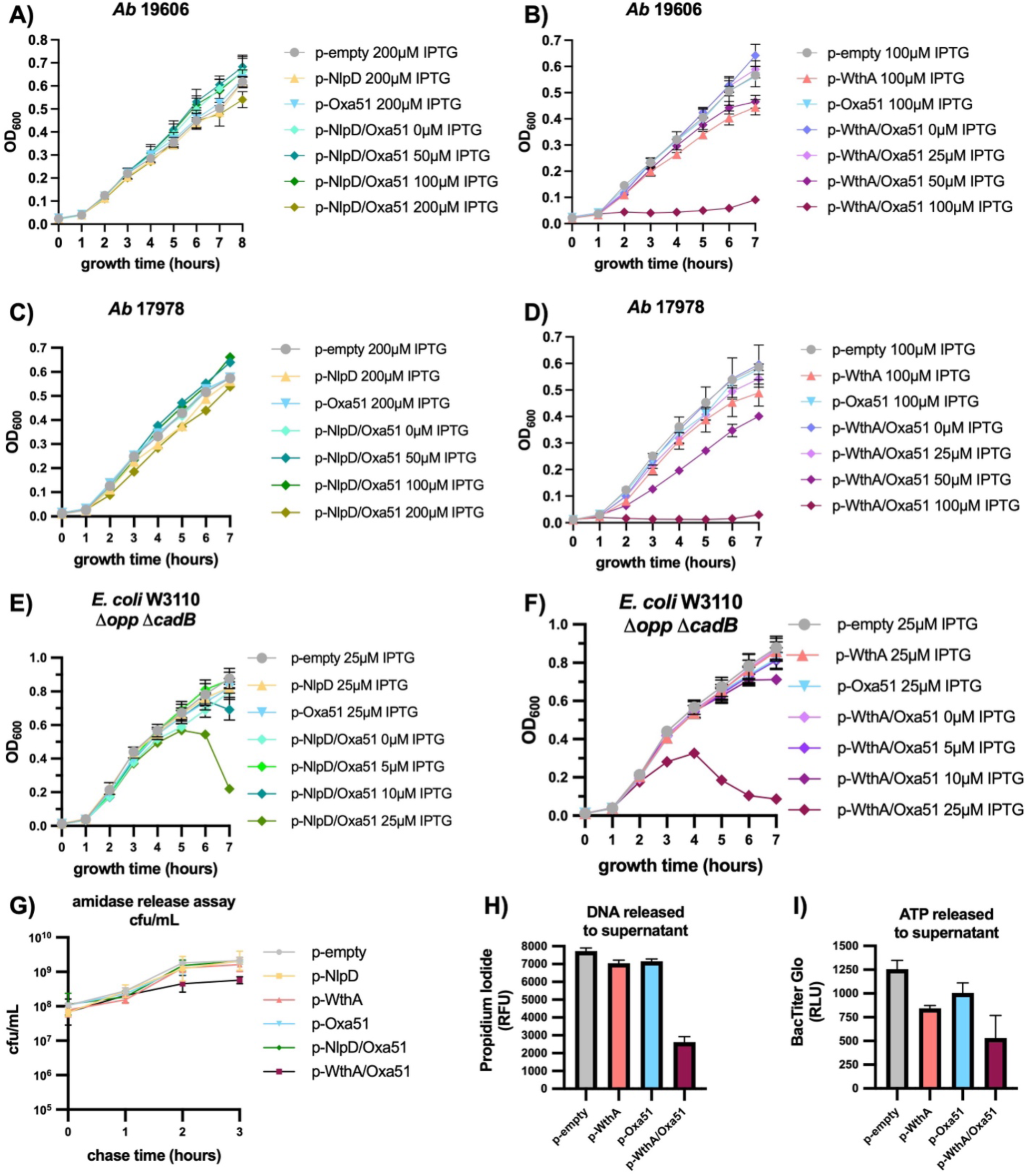
Impact on growth during WthA and NlpD coexpression with Oxa51. (A-F) Growth curves of single or coexpressions of Oxa51 and putative amidase activators NlpD (A,C,E) or WthA (B,D,F) expressed in *A. baumannii* ATCC 19606 (A-B), *A. baumannii* ATCC 17978 (C-D), or *E. coli* W3110 Δ*opp* Δ*cadB* (also called AmpG only strain, C-D). All growth curves are biological triplicates grown in LB with the indicated concentrations of IPTG. (G) Colony forming units determined during the three-hour chase of Amidase Product Release Assays in **Fig. 6D**. *E. coli* W3110 Δ*opp* Δ*cadB* strain expressing proteins on indicated plasmids with 5 µM IPTG. (H-I) Controls for release of cytoplasmic contents into the supernatant at the end-point of the Amidase Product Release Assays in **Fig. 6D**. Supernatants were test for DNA with propidium iodide fluorescence (H) and for ATP with BacTiter Glo luciferase luminescence (I) in technical duplicate of biological triplicates. Error bars for all experiments indicate standard deviation and are not shown when smaller than the size of the symbol.

**Figure S11.**
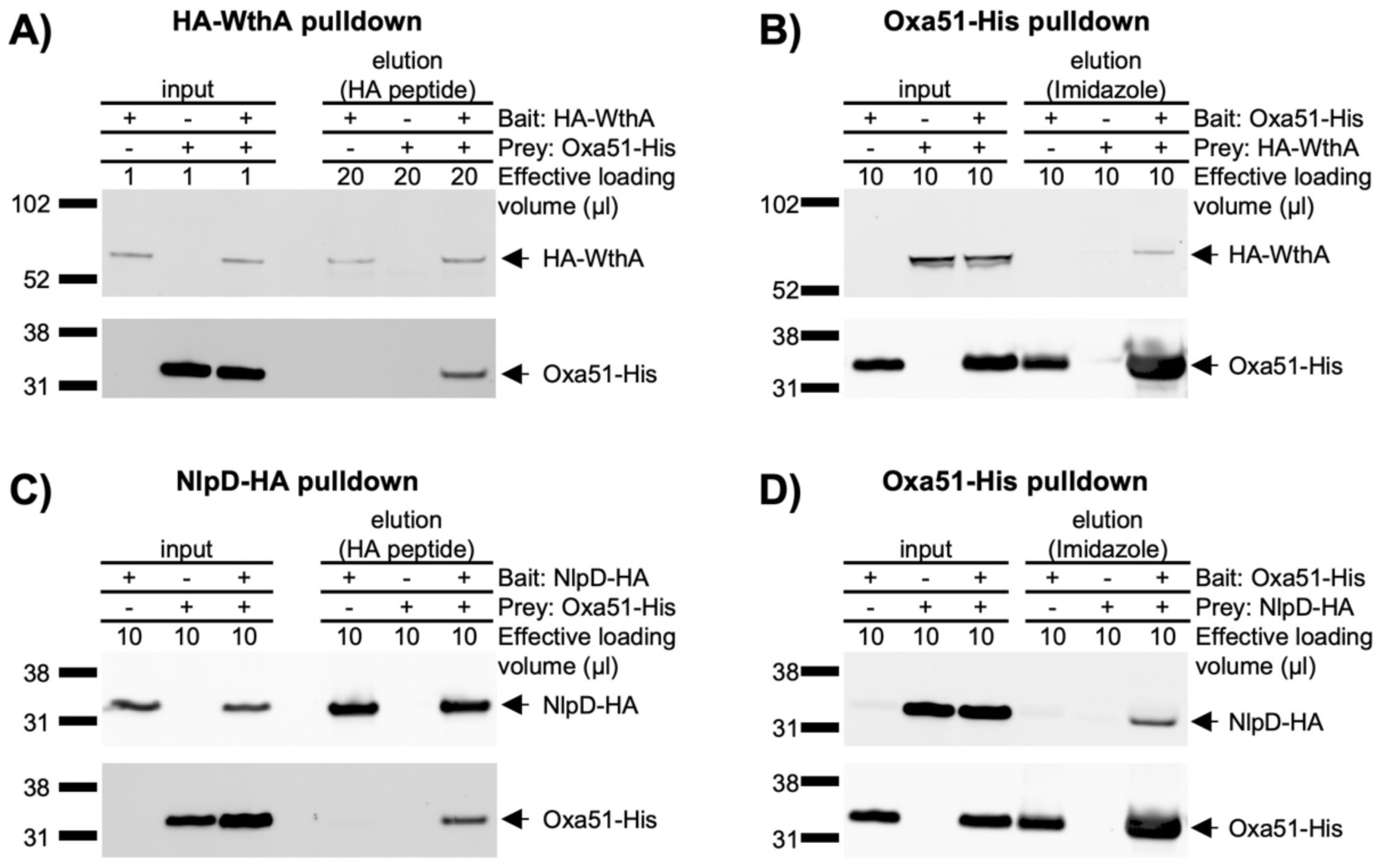
Evidence of interactions between Oxa51 and WthA or NlpD. Anti-HA (top of each panel) and anti-His (bottom of each panel) immunoblots of lysate inputs and protein elutions from indicated pull downs. (A-B) Anti-HA immunoprecipitation (A) and nickel affinity His pull down (B) of protein lysates from *A. baumannii* ATCC 17978 cells expressing HA-WthA and Oxa51-His. (C-D) Anti-HA immunoprecipitation (C) and nickel affinity His pull down (D) of protein lysates from *A. baumannii* ATCC 17978 cells expressing NlpD-HA and Oxa51-His. All pull-down experiments are representative of biological triplicates. Loading volumes are indicated.

**Figure S12.**
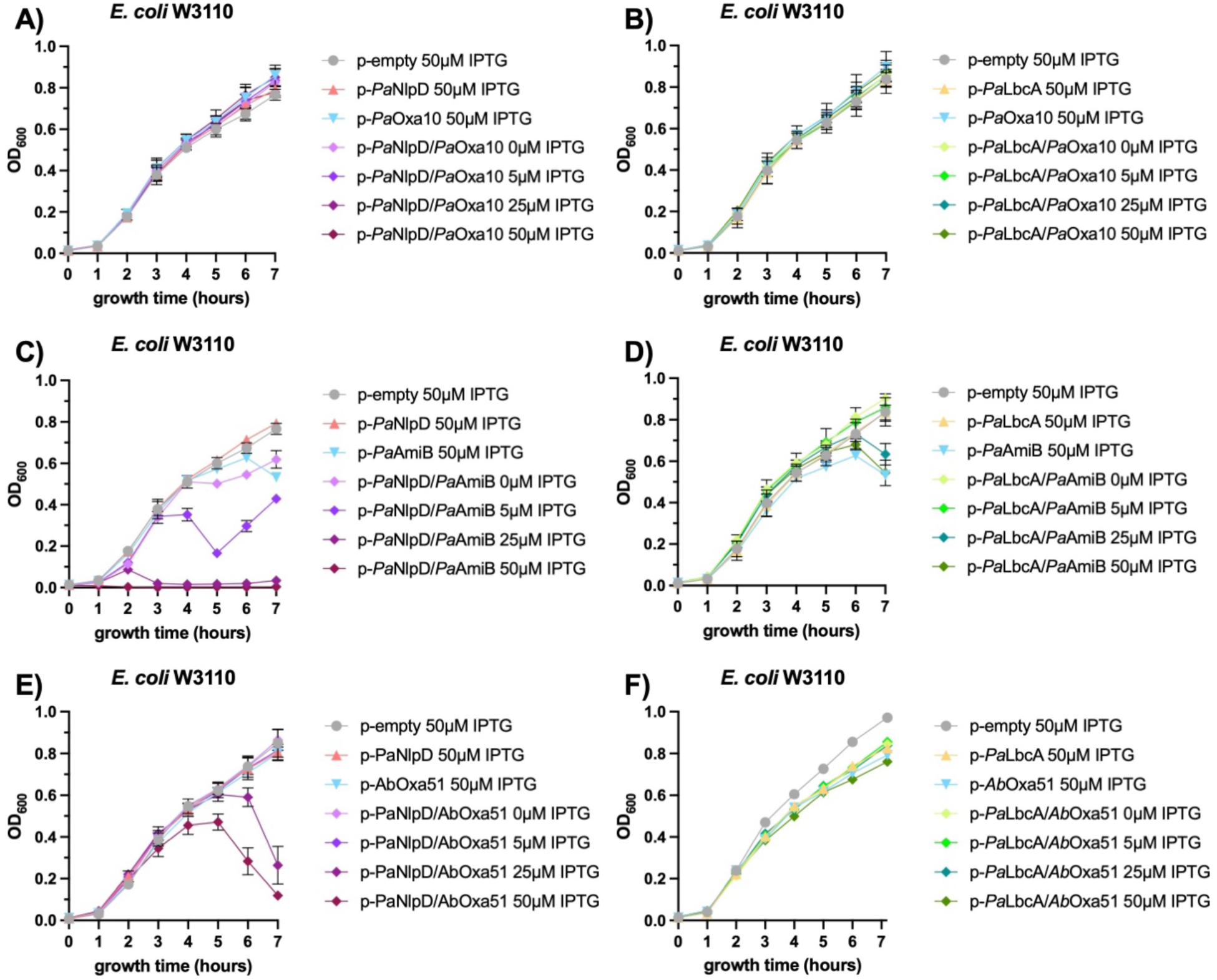
Toxicity screen to test *P. aeruginosa* homologs for amidase activation. Growth curves in biological triplicate of *E. coli* W3110 expressing plasmids with possible amidases, *Pa*Oxa10 (A-B), *Pa*AmiB (C-D), and *Ab*Oxa51 (E-F) alone or in combination with possible amidase activators *Pa*NlpD (A,C,E) or *Pa*LbcA (B,D,F). All growth curves were grown in LB and contained the indicated concentrations of IPTG. Error bars indicate standard deviation and are not shown when smaller than the size of the symbol.

---

**Table S1.** Antibiotic sensitivity profile of *wthA* mutants indicates low PBP activity and uncrosslinked D-Ala-D-Ala.

| Antibiotic | 19606 MIC<br>(Avg ± SD) | $\Delta wthA$ MIC<br>(Avg ± SD) | Fold change | Target | Interpretation |
| --- | --- | --- | --- | --- | --- |
| Rifampicin | 0.75 ± 0.00 | 0.104 ± 0.018 | -7.2 | RNA pol/ hydrophobic | OM disruption |
| Fosfomycin | 85.3 ± 18.5 | 69.3 ± 24.4 | -1.2 | Lipid II synthesis | no impact |
| Vancomycin | 74.7 ± 18.5 | >256 | >3.4 | D-Ala-D-Ala | Uncrosslinked<br>D-Ala-D-Ala |
| Ceftazidime-avibactam | 1.83 ± 0.29 | 0.46 ± 0.07 | -4.0 | PG transpeptidases<br>(PBP2/3/1A/1B) | decreased<br>crosslinks |
| Aztreonam | 12.0 ± 0.0 | 4.67 ± 1.15 | -2.6 | PBP3 | cell division defect |
| Bacitracin | 42.7 ± 9.2 | 32 ± 13.9 | -1.3 | UndP | no impact |

